# Adaptive disinhibitory gating by VIP interneurons permits associative learning

**DOI:** 10.1101/443614

**Authors:** Sabine Krabbe, Enrica Paradiso, Simon D’Aquin, Yael Bitterman, Chun Xu, Keisuke Yonehara, Milica Markovic, Jan Gründemann, Francesco Ferraguti, Andreas Lüthi

**Author notes:** These authors contributed equally.

## Abstract

Learning drives behavioral adaptations necessary for survival. While plasticity of excitatory projection neurons during associative learning is studied extensively, little is known about the contributions of local interneurons. Using fear conditioning as a model for associative learning, we find that behaviorally relevant, salient stimuli cause learning by tapping into a local microcircuit consisting of precisely connected subtypes of inhibitory interneurons. By employing calcium imaging and optogenetics, we demonstrate that vasoactive intestinal peptide (VIP)-expressing interneurons in the basolateral amygdala are activated by aversive events and provide an instructive disinhibitory signal for associative learning. Notably, VIP interneuron responses are plastic and shift from the instructive to the predictive cue upon memory formation. We describe a novel form of adaptive disinhibitory gating by VIP interneurons that allows to discriminate unexpected, important from irrelevant information, and might be a general dynamic circuit motif to trigger stimulus-specific learning, thereby ensuring appropriate behavioral adaptations to salient events.

## INTRODUCTION

Associative learning allows an organism to link environmental cues with their motivational and emotional significance. Mechanisms of memory formation are critically shaped by dynamic changes in the balance of excitatory and inhibitory neuronal circuit elements (Froemke, 2015). Although local inhibitory interneurons only represent a minority of the cells in cortical areas, they can tightly regulate the activity of excitatory projection neurons (PNs) in a spatially and temporally precise manner (Kepecs and Fishell, 2014). However, to date, it remains largely unknown how distinct interneuron subtypes contribute to memory formation and plasticity *in vivo*, and how they control the transformation of sensory information to change the appropriate behavioral output upon learning.

Learning is often driven by unexpected positive or negative experiences in an ever-changing environment. Auditory fear conditioning in rodents, in which an initially neutral tone (conditioned stimulus, CS) is paired with an aversive stimulus (unconditioned stimulus, US), is one of the most powerful model systems to investigate the neuronal mechanisms of this associative memory formation (Fanselow and Poulos, 2005; LeDoux, 2000; Tovote et al., 2015). The US, typically a mild foot shock, can be regarded as teaching signal, instructing neuronal plasticity to the auditory cue (Ozawa and Johansen, 2018). However, repeated predicted presentations of a US will reduce its salience. Accordingly, neuronal activity induced by instructive signals can be modulated by memory formation, and responses decrease upon learning of the CS-US contingency (McNally et al., 2011;Ozawa and Johansen, 2018). Yet, it is unclear how this mechanism of expectation-dependent modulation of the representation of salient cues is shaped by local microcircuits.

The basolateral amygdala (BLA) is a major site of synaptic plasticity during associative fear learning (Fanselow and Poulos, 2005; LeDoux, 2000; Tovote et al., 2015). Anatomically, the BLA represents a cortex-like structure, consisting of a majority of excitatory PNs and only about 20% inhibitory interneurons (Krabbe et al., 2018). Two major subclasses of BLA GABAergic interneurons expressing parvalbumin (PV) and somatostatin (SOM) have previously been identified to exert powerful inhibitory control over PNs, with PV cells preferentially targeting their perisomatic region and SOM cells inhibiting their distal dendrites (Wolff et al., 2014). Release from this inhibition and depolarization of BLA PNs during instructive cues has been shown to regulate aversive learning (Johansen et al., 2010a; Krabbe et al., 2018; Wolff et al., 2014), but the source and spatial pattern of this disinhibition is so far unknown.

Studies in the neocortex identified an independent class of interneurons, vasoactive intestinal peptide (VIP)-expressing cells, as upstream modulator of both PV and SOM interneurons, building a local microcircuit that can lead to disinhibition of PNs (Lee et al., 2013; Pfeffer et al., 2013; Pi et al., 2013). A similar layout of preferential interneuron targeting by VIP cells has been proposed for the BLA (Muller et al., 2003; Rhomberg et al., 2018). Furthermore, cortical VIP interneurons can be activated by salient stimuli such as appetitive and aversive cues (Pi et al., 2013). However, the behavioral relevance of this activity pattern has not been demonstrated yet. Based on these observations, we hypothesized that a disinhibitory VIP → PV and/or SOM → PN circuit motif in the BLA could be involved in associative learning instructed by salient aversive stimuli.

To test this, we used a combination of deep brain imaging of BLA interneurons and PNs, as well as optogenetic manipulations in freely behaving mice. In this study, we report that VIP BLA interneurons are strongly activated by aversive cues during associative fear learning and thereby provide a mandatory signal for memory formation. Similar to cortex, VIP BLA interneurons preferentially target SOM and PV interneurons and thus promote PN depolarization during aversive stimuli. We further show that this inhibitory gating, likely dominated by dendritic disinhibition, permits BLA PN plasticity to predictive cues. Remarkably, VIP BLA interneurons exhibit a teaching signal pattern during fear learning, and shift their activity from the aversive to the predictive cue upon formation of the association. Based on our data, we propose that this novel form of adaptive disinhibitory gating represents a key mechanism for the computation of unexpected, meaningful events in local microcircuits, and allows for plastic changes in excitatory principal circuit elements to ensure behavioral adaptations to salient environmental cues.

## RESULTS

### Salient aversive cues activate VIP BLA interneurons

To investigate the functional role of VIP interneurons for associative learning instructed by aversive events, we employed a deep brain Ca^2+^ imaging approach combined with an ultra-light head-mountable miniaturized microscope (Ghosh et al., 2011; Grewe et al., 2017; Gründemann et al., 2018). Cre-dependent, virally-mediated expression of GCaMP6s (Chen et al., 2013) and the implantation of gradient-index (GRIN) lenses in the BLA of VIP-cre mice allowed VIP interneuron specific Ca^2+^ imaging in freely moving mice at single cell resolution (**Figures 1A-D,F and S1A-C**). Mice with head-mounted miniature microscopes underwent a discriminative fear conditioning paradigm, in which five presentations of the auditory CS+ were paired with an aversive US, while intermingled CS presentations were used as control tones (**Figures 1E and S1D**). A large fraction of VIP BLA interneurons displayed strong activation upon US application (76% of n=170 cells from N=7 mice; **Figure 1G-I**). While a substantial proportion of cells was also mildly activated by the predictive CS+ (64%), only a small subset responded to the non-reinforced CS (24%; **Figure 1H-I**). Notably, US responses of individual VIP interneurons decreased across repeated trials, while learning, measured by freezing during the predictive CS+, progressed (**Figure 1J**).

**Figure 1.**
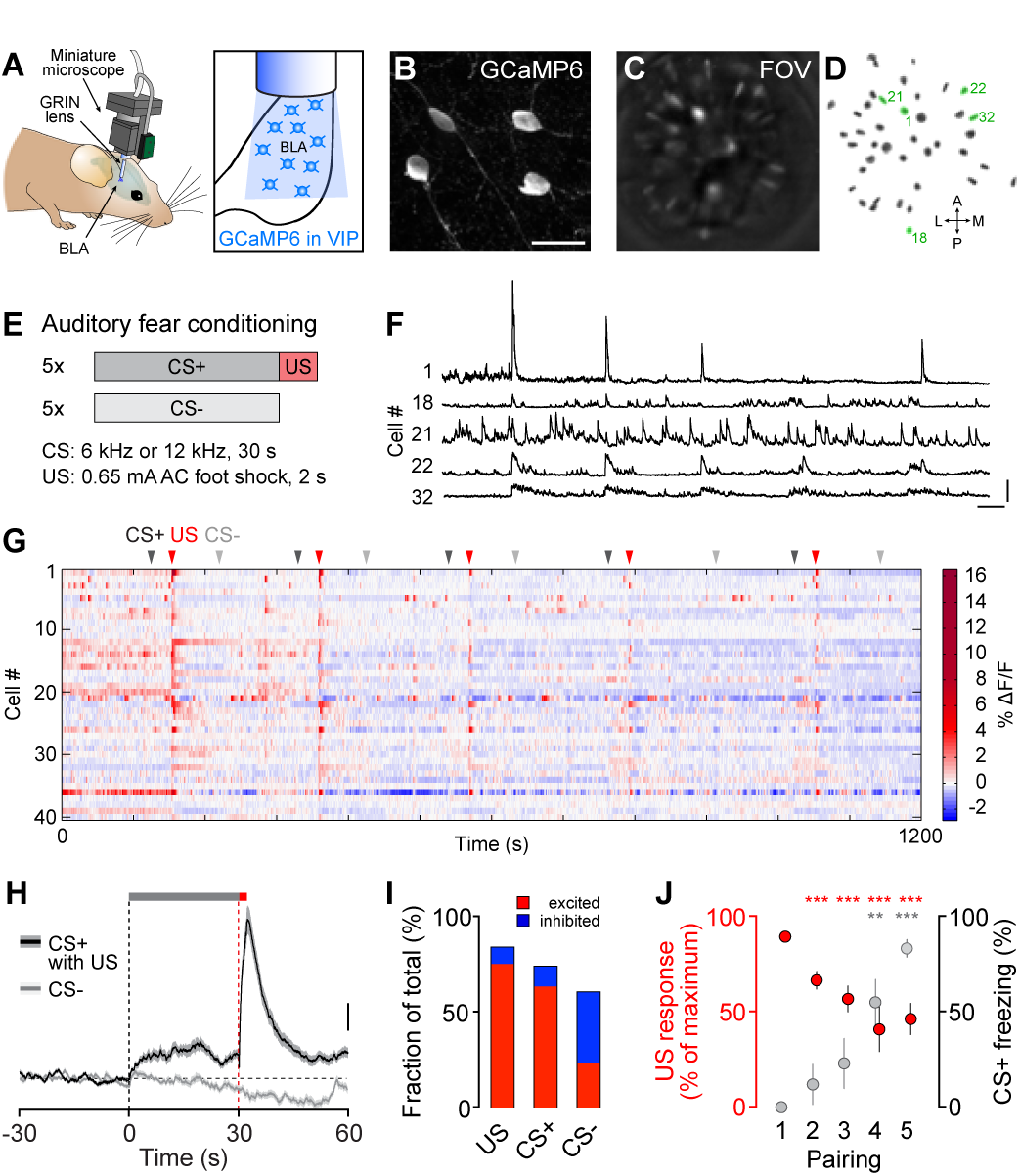
Aversive foot shocks activate VIP BLA interneurons during fear learning. **(A)** Schematic illustrating miniature microscope and implanted gradient-index (GRIN) lens for deep brain Ca^2+^ imaging of BLA interneurons in freely behaving mice. **(B)** Representative confocal example image of cre-dependent GCaMP6 expression in VIP interneurons in the BLA of VIP-cre mice. Scale bar, 20 μm. **(C)** Example field of view (FOV) in the BLA through the implanted GRIN lens using the miniature microscope five weeks after cre-dependent GCaMP6 injection in VIP-cre mice. **(D)** Spatial filters of identified VIP BLA interneurons in the same mouse (n=40 cells). **(E)** Schematic showing discriminative auditory fear conditioning paradigm used for miniature microscope imaging. CS+ and US pairings were presented alternating with the CS-. **(F)** Ca^2+^ traces from the entire behavioral session (20 min) from five example VIP BLA interneurons (numbers correspond to labelled spatial filters in D). Scale bars, 10% ΔF/F, 60 s. **(G)** Activity map of all identified cells from the example mouse across the entire fear conditioning session. Arrowheads indicate onset of CS+ and US as well as intermingled CS-. **(H)** CS and US responses from all recorded VIP BLA interneurons (n=170 cells from N=7 mice) averaged across all five trials (traces represent mean in black and s.e.m. in lighter gray). Scale bar, 0.2% ΔF/F. **(I)** Percentage of VIP BLA interneurons with significantly increased or decreased Ca^2+^ responses during distinct stimulus presentations (n=170) based on averages across all five trials. **(J)** US responses decrease with repeated pairings of CS+ and US in VIP BLA interneurons, while learning measured by freezing during the CS+ increases in the same mice (US response: Kruskal-Wallis test, H=149, P<0.0001; Dunn’s multiple comparisons test, US1 vs US2, P<0.0001; vs US3, P<0.0001; vs US4, P<0.0001; vs US5, P<0.0001; n=130 first US excited cells from N=7 mice; CS freezing: Kruskal-Wallis test, H=22.17, P<0.001; Dunn’s multiple comparisons test, CS1 vs CS4, P<0.01; vs CS5, P<0.001; N=7). Data in H and J is shown as mean and s.e.m. ** P<0.01, *** P<0.001. Details of statistical analysis are listed in Table S3.

Previous studies using electrophysiological single unit recordings reported heterogeneous response patterns to aversive stimuli in PV and SOM BLA interneurons (Bienvenu et al., 2012; Wolff et al., 2014). To monitor the activity of larger populations of PV and SOM BLA interneurons within single animals, we applied our deep brain imaging approach during associative fear learning to these inhibitory subtypes by viral expression of GCaMP in the BLA of PV-cre and SOM-cre mice (**Figure S1E-H**). CS+ and US response patterns of PV interneurons were similar to those of VIP interneurons, although the fraction of US inhibited cells was slightly larger (20% of n=46 cells from N=4 mice; **Figure S1F, I-K**). In contrast, SOM interneurons exhibited significantly different response profiles with larger proportions of inhibited cells during both the aversive US (34% of n=152 cells from N=5 mice) and the predictive CS+ (38%; **Figure S1H, I-K**). Overall, all three BLA interneuron subtypes showed large response heterogeneity to both predictive, instructive and neutral sensory stimuli within individual mice, yet VIP interneurons were found to exhibit the strongest and most uniform activity to aversive cues, while a large fraction of the PN dendrite-targeting SOM interneurons was simultaneously inhibited.

### A unique position of VIP interneurons in a highly interconnected BLA microcircuit

In light of the distinct response patterns in these three non-overlapping BLA interneuron subpopulations, we next analyzed their presynaptic long-range connectivity which might drive the stimulus-specific activity. Using monosynaptic rabies tracing (Callaway and Luo, 2015), we mapped whole-brain presynaptic inputs to VIP, PV and SOM BLA interneurons. To this end, we injected a cre-dependent AAV encoding the TVA receptor and rabies glycoprotein unilaterally into the BLA of VIP-cre, PV-cre or SOM-cre mice, followed by the injection of rabiesΔG two weeks later (**Figures 2A and S2**). VIP interneurons received major inputs from auditory areas in cortex and thalamus, as well as from brain regions involved in aversive signaling such as the dorsal midline and intralaminar thalamus as well as the insular cortex (Lanuza et al., 2008; 2004; Sengupta and McNally, 2014) (**Figures 2B-F and S3A-B**). We further observed strong rabies labeling in the basal forebrain, which in part comprised cholinergic basal forebrain neurons (**Figures 2C and S3C**). High numbers of cells monosynaptically connected to VIP interneurons were also found in piriform cortex, entorhinal and perirhinal cortex, as well as the ventral hippocampus (**Figures 2G and S3A-B**).

**Figure 2.**
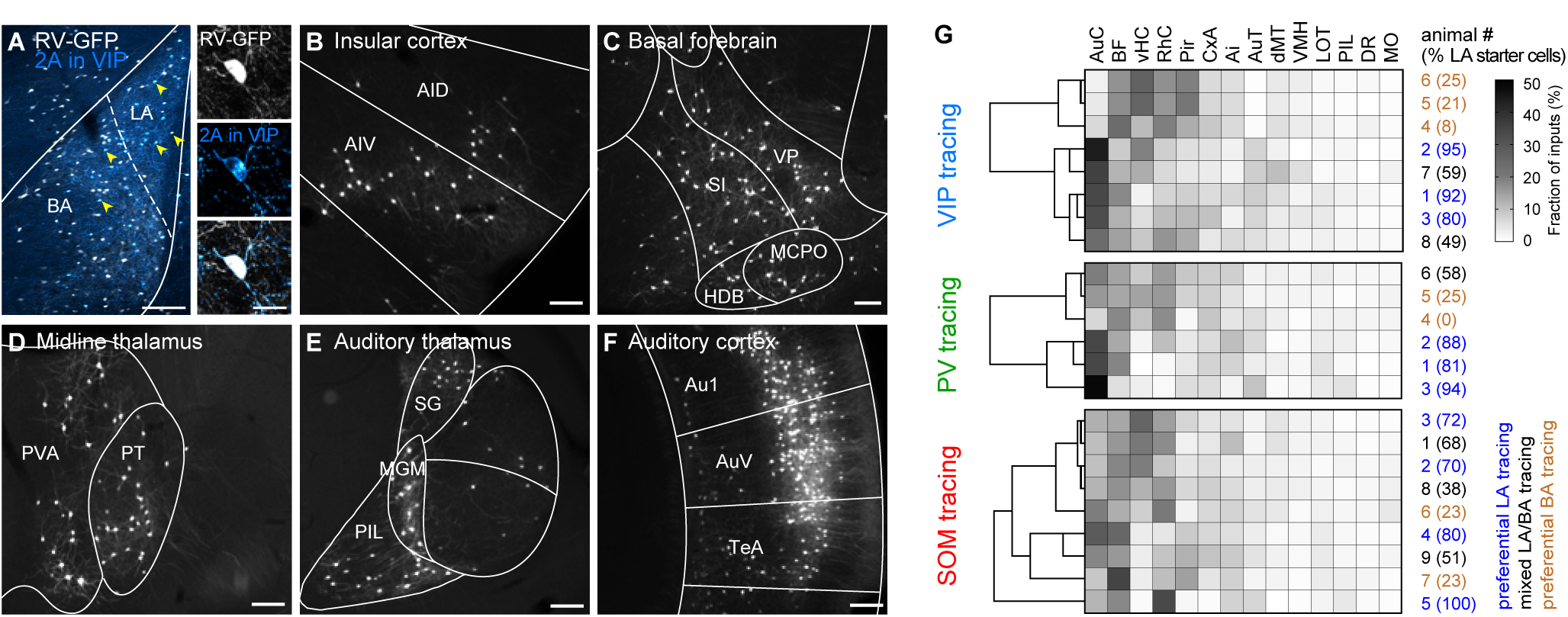
Long-range connectivity of distinct BLA interneuron subtypes. **(A)** Representative example image of 2A peptide and rabies-GFP (RV-GFP) expression in VIP interneurons in the BLA of VIP-cre mice. Yellow arrowheads point to identified starter cells expressing both TVA950-2A-CVS11G and RV-GFP. LA, lateral amygdala; BA, basal amygdala. Scale bar, 200 μm. High magnification shows an example starter cell. Scale bar, 20 μm. **(B-F)** Monosynaptic inputs to BLA VIP interneurons from (B) insular cortex, (C) basal forebrain, (D) dorsal midline thalamus, (E) auditory thalamus and (F) auditory cortex. For anatomical abbreviations, see methods. Scale bar, 200 μm. **(G)** Dendrograms and heatmaps representing hierarchical clustering of VIP, PV and SOM BLA interneuron tracings based on fraction of inputs (VIP, N=8 mice; PV, N=6; SOM, N=9; cluster method, between groups linkage; measure, squared Euclidean distance). Each matrix row depicts a single mouse. Example tracing in A-F is from mouse #8 in VIP-cre heatmap. For anatomical abbreviations, see methods or Figure S3. Results from rabies tracings are summarized in Table S1.

In accordance with our *in vivo* imaging data, PV BLA interneurons showed a very similar input pattern compared to VIP cells (**Figures 2G and S3A-B**). Moreover, hierarchical cluster analysis revealed that VIP-cre and PV-cre mice with preferential lateral (LA) or basal (BA) amygdala tracing injections clustered together, with sensory brain areas mainly targeting the respective LA interneurons, while higher order processing areas such as the ventral hippocampus or rhinal cortices mainly targeted BA populations (**Figure 2G**). In contrast, we found no differential pattern of presynaptic inputs to SOM interneurons in the LA or in the BA (**Figures 2G and S3A-B**). This suggests that BLA SOM interneurons receive a different array of afferent innervation compared to VIP and PV interneurons, which might contribute to the differential activity patterns observed during associative fear learning.

We next examined the local connectivity between distinct BLA interneuron subtypes and neighboring PNs. First, we expressed the excitatory opsin ChR2 specifically in VIP, PV or SOM interneurons of the BLA. Using whole-cell patch-clamp experiments in acute brain slices, we recorded inhibitory postsynaptic currents (IPSCs) in neighboring PNs in response to brief light stimulation (**Figure 3A-C**). Both PV and SOM interneuron network photostimulation induced strong short-latency IPSCs in almost all recorded BLA PNs (97% of n=35 cells from N=6 mice for PV, 100% of n=33 cells from N=7 mice for SOM; **Figures 3A-B and Table S2**). In contrast, less than half of the recorded PNs received inhibition from VIP cells (49% of n=72 cells from N=13 mice; **Figure 3C-D**). Moreover, these sparse inhibitory inputs were significantly weaker compared to PV-and SOM-induced IPSCs (**Figure 3E-F**).

**Figure 3.**
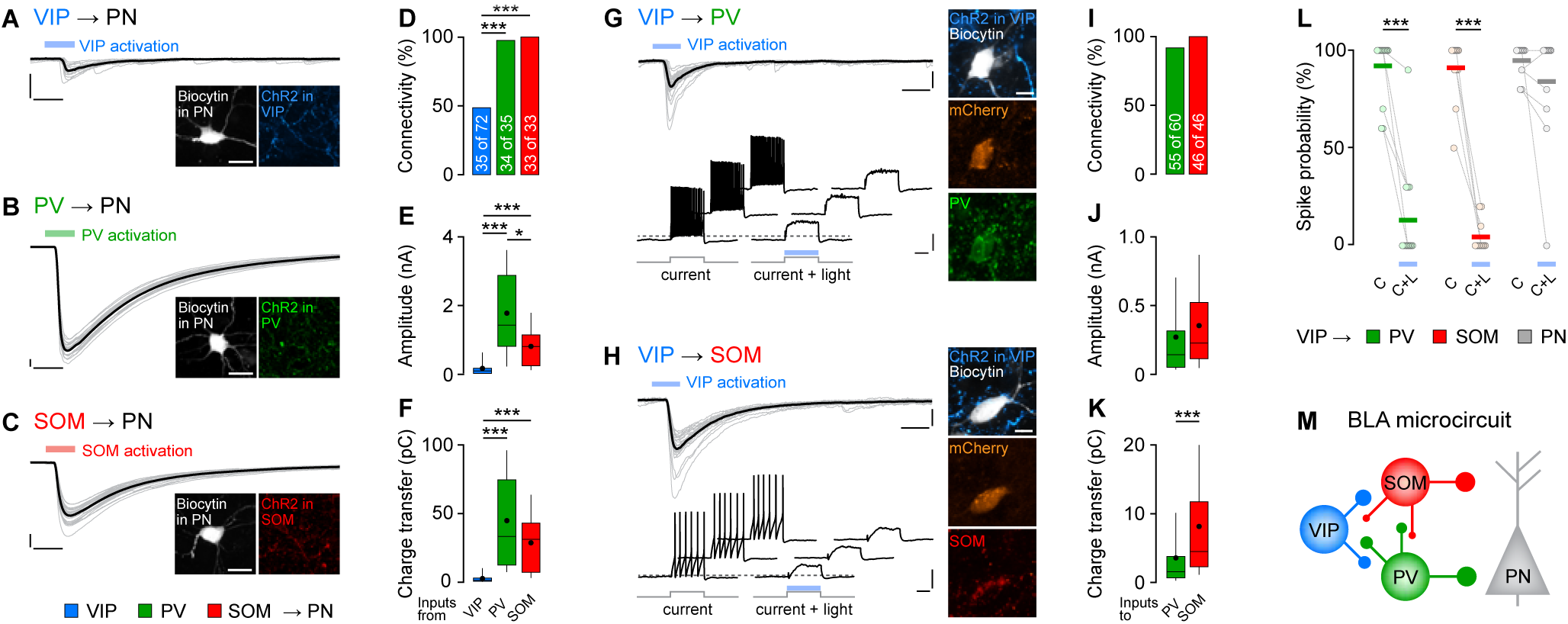
VIP BLA interneurons preferentially target other interneuron subtypes over principal cells. **(A-C)** Inhibitory post-synaptic currents (IPSCs) evoked in BLA PNs by brief photostimulation (color-coded bars) of (A) VIP, (B) PV and (C) SOM interneuron networks expressing ChR2. Scale bars, 200 pA, 10 ms. Corresponding confocal images show biocytin-filled recorded PNs. Scale bars, 20 μm. **(D)** Connectivity is significantly higher from PV and SOM BLA interneurons onto PNs compared to VIP interneurons (VIP, 48.6%, n=72 cells from N=13 mice; PV, 97.1%, n=35, N=6; SOM, 100%, n=33, N=7; Pearson’s X^2^ test P<0.0001; Fisher’s exact post-hoc test VIP vs PV, P<0.0001; VIP vs SOM: P<0.0001). **(E-F)** Sparse inputs from VIP interneurons exhibit significantly weaker (E) amplitudes (Kruskal-Wallis test, H=42.79, P<0.0001; Dunn’s multiple comparisons test, VIP vs PV, P<0.0001, VIP vs SOM: P<0.001, PV vs SOM: P<0.05; VIP, n=22 cells, PV, n=34, SOM, n=33) and (F) lower charge transfer compared to PV and SOM stimulation (Kruskal-Wallis test, H=41.18, P<0.0001; Dunn’s multiple comparisons test, VIP vs PV, P<0.0001, VIP vs SOM: P<0.0001). **(G)** Top, short-latency IPSCs in PV BLA interneurons upon VIP network activation (blue bar). Scale bars, 100 pA, 10 ms. Right, corresponding confocal image confirming PV expression. Scale bar, 20 μm. Bottom, VIP BLA activation can reliably suppress spiking in PV cells. Dashed line, −50 mV. Scale bars, 20 mV, 200 ms. **(H)** Top, example IPSCs from a SOM BLA interneuron upon VIP BLA photostimulation with corresponding confocal image confirming SOM expression. Scale bars, 100 pA, 10 ms; 20 μm. Bottom, spike-suppression in a SOM BLA interneurons upon VIP activation. Dashed line, −65 mV. Scale bars, 20 mV, 200 ms. **(I)** High connectivity from VIP BLA interneurons to PV and SOM interneurons (PV, 91.7%, n=60, N=7; SOM, 100%, n=46, N=5). **(J-K)** Average (J) amplitude and (K) charge transfer (Mann-Whitney U test, P<0.001; PV, n=41; SOM, n=34) of light-induced IPSCs from VIP to PV and SOM BLA interneurons. **(L)** Reliable spike suppression upon VIP network activation in PV and SOM interneurons, but not BLA PNs (PV, Wilcoxon matched-pairs signed rank test, P<0.001, n=14; SOM, Wilcoxon matched-pairs signed rank test, P<0.001, n=12). **(M)** Proposed BLA microcircuit based on *ex vivo* connectivity assays with reciprocal interneuron connectivity with variable strength (see also Figure S4). Individual IPSC traces from one cell are gray, corresponding average black in panels (A-C) and (G-H). Box-whisker plots show median values and 25^th^/75^th^ percentiles with 10^th^ to 90^th^ percentile whiskers, dots indicate the mean. Circles in panel L represent individual data points, horizontal lines the mean. * P<0.05, ** P<0.01, *** P<0.001. All results from slice electrophysiology analysis are summarized in Table S2, details of statistical analysis are listed in Table S3.

Next, we assessed interneuron interconnectivity and recorded VIP inputs to PV or SOM cells using a dual-transgenic mouse line approach, in which we expressed ChR2 cre-dependently in VIP interneurons, and the marker mCherry flp dependently in PV or SOM interneurons of the BLA. We reliably observed light-induced ABAergic inputs to PV and SOM interneurons when activating the VIP network (92% of n=60 cells from N=7 mice for PV, 100% of n=46 cells from N=5 mice for SOM; **Figures 3G-I and S4B-E**), with slightly stronger currents in SOM compared to PV cells (**Figure 3J-K**). Importantly, spiking activity in PV and SOM BLA interneurons, but not PNs, could be robustly suppressed by simultaneous photostimulation of VIP interneurons (**Figure 3G-H, L**).

Circuit mapping approaches in cortex recently suggested reciprocal connectivity between distinct subtypes of inhibitory interneurons (Dipoppa et al., 2018; Jiang et al., 2015; Pfeffer et al., 2013). Therefore, we additionally examined the level of interconnectivity between the distinct BLA interneuron populations. Indeed, we found that VIP, PV and SOM BLA interneurons are reciprocally connected (**Figures 3M, S4F-O and Table S2**). However, VIP interneurons have a unique position in this BLA microcircuit as they almost exclusively inhibit other interneuron subtypes, while PV and SOM interneurons exert their strongest inhibitory control over PNs.

### VIP BLA interneurons control projection neuron plasticity and learning by disinhibition

In this exquisitely organized BLA microcircuit, VIP interneurons are in an ideal position to gate information flow to PNs during the aversive US by releasing them from the strong inhibition provided by SOM and PV interneurons (Letzkus et al., 2015; Wolff et al., 2014). This suggests that US-induced VIP activity would promote PN depolarization and thus ultimately support associative learning (Johansen et al., 2010a; Sengupta et al., 2018). To test this hypothesis, we combined simultaneous deep brain Ca^2+^ imaging and optogenetic manipulations in freely behaving mice. We first tested the efficiency of optogenetic manipulations on US-evoked responses in VIP BLA interneurons by expressing GCaMP together with the inhibitory opsin ArchT in the BLA of VIP-cre mice (**Figure 4A-B**). After confirming specific expression of ArchT in VIP interneurons and functionality of the construct (**Figure S5A-E**), we verified in acute brain slices that the opsin was reliably activated by yellow, and not by the low-intensity blue light used for Ca^2+^ imaging (**Figure S5F-G**). Conversely, yellow light used for optogenetic manipulation did not affect the GCaMP signal in control mice, but significantly reduced the Ca^2+^ signal in cells with ArchT co-expression under baseline conditions *in vivo* (**Figure S5H-K**).

**Figure 4.**
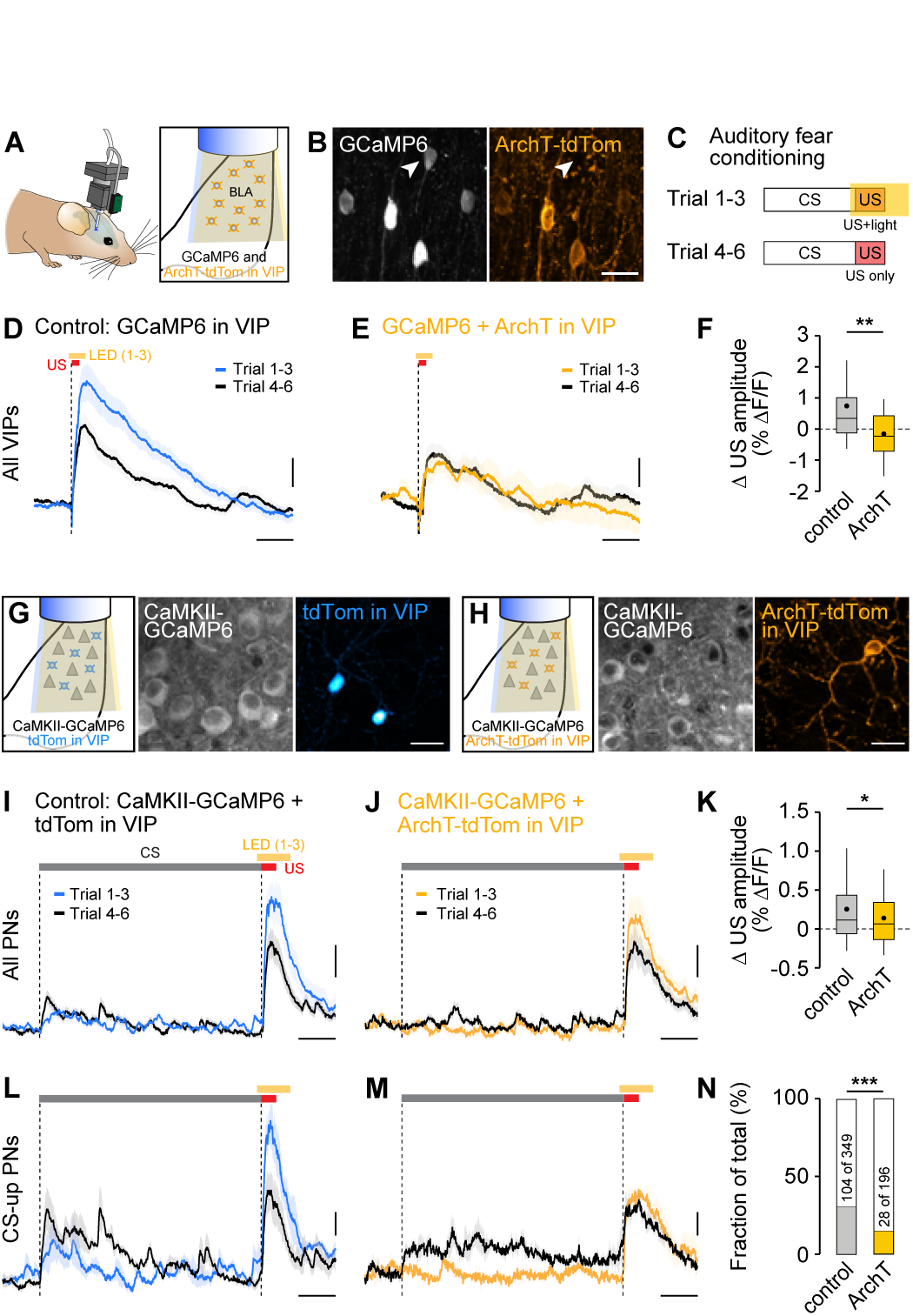
VIP BLA interneuron activation disinhibits projection neurons. **(A)** Approach for combined deep brain Ca^2+^ imaging and optogenetic manipulation using the nVoke miniature microscope system. Blue light (448 nm) for GCaMP imaging and yellow light (590 nm) for inhibition of cellular activity with ArchT are transmitted through the objective of the microscope and the implanted GRIN lens. **(B)** Selective expression of GCaMP6 and ArchT-tdTomato in VIP BLA interneurons of VIP-cre mice. Few cells expressed GCaMP, but not ArchT (arrowhead). Scale bar, 20 μm. **(C)** Schematic illustrating behavioral protocol for nVoke fear conditioning experiments. Six pairings of CS and US were applied, yellow light for ArchT activation was administered during the US of the first three trials, while the last three trials served as internal control. **(D)** Average responses to the US during light-modulated (blue trace) and no-light trials (gray trace) in VIP BLA interneurons in a control group of mice expressing only GCaMP6 (red bar starting from dashed line marks the US, yellow bar the light duration in trials 1-3). Consistent with previous results, the first three pairings induce stronger US responses in VIP interneurons compared to the last three trials (n=133 cells from N=4 mice). Scale bar, 0.5% ΔF/F, 10 s. **(E)** Average US responses to light-modulated (yellow) and no-light trials (gray) in VIP BLA interneurons expressing GCaMP6 and ArchT-tdTomato. US activation is not completely abolished, but drastically decreased by simultaneous ArchT activation with yellow light (n=25, N=1). Scale bar, 0.5% ΔF/F, 10 s. **(F)** Difference in maximum US responses between light and no-light trials is significantly greater in cells from control mice compared to cells with ArchT co-expression (Mann-Whitney U test, P<0.01; control, n=133; ArchT, n=25). **(G-K)** Combined deep brain imaging of BLA PNs and optogenetic manipulation of VIP interneurons. **(G)** Strategy for control mice, expressing GCaMP6 in BLA projections neurons with tdTomato in VIP interneurons. Scale bar, 20 μm. **(H)** Combined expression of GCaMP6 in BLA PNs and ArchT-tdTomato in VIP interneurons for optogenetic manipulation. Scale bar, 20 μm. **(I)** Average CS and US responses to light-modulated (blue) and no-light trials (gray) in BLA PNs in the control group expressing tdTomato in VIP BLA interneurons. Similar to VIP interneurons, the first three trials induce stronger US responses in BLA PNs compared to the last three trials (n=349, N=6). **(J)** Average CS and US responses in BLA PNs during trials with simultaneous VIP inhibition using ArchT (yellow) and no-light control trials (gray; n=196, N=4). **(K)** Difference in maximum US responses between light and no-light trials is significantly smaller in PNs with ArchT-dependent VIP modulation compared to the control group (Mann-Whitney U test, P<0.05; control, n=349; ArchT, n=196). **(L)** Response profiles of CS-up cells in BLA PNs of control mice during US-light (blue) and no-light trials (gray; n=104, N=6). **(M)** Response profiles of CS-up cells in BLA PNs during trials with (yellow) and without (gray) VIP inhibition during the US (n=28, N=4). **(N)** VIP inhibition during the aversive US significantly reduces the fraction of CS-up cells in the BLA PN population (Pearson’s X^2^ test, P<0.0001; control, n=349; ArchT, n=196). Scale bar I-J, L-M, 0.1% ΔF/F, 5 s. Box-whisker plots show median values and 25^th^/75^th^ percentiles with 10^th^ to 90^th^ percentile whiskers, dots additionally indicate the mean. * P<0.05, ** P<0.01, *** P<0.001. Details of statistical analysis are listed in Table S3.

Freely behaving mice with head-mounted miniaturized microscopes were exposed to an auditory fear conditioning paradigm with six CS-US pairings. Only for the first half of the trials the US was accompanied by yellow light for ArchT activation, while the last trials were used as internal control (**Figure 4C**). Consistent with the previously observed decrease in VIP US responses across repeated CS-US pairings (**Figure 1J**), we found that in control mice expressing GCaMP alone, the first three trials induced stronger US activity compared to the last three trials (**Figure 4D**). In contrast, simultaneous optogenetic activation of ArchT significantly decreased US-evoked Ca^2+^ responses within the VIP BLA population during auditory fear conditioning (**Figure 4E-F**),demonstrating a high efficiency of our optogenetic silencing approach during the aversive US.

To examine the impact of VIP interneuron activity during the US on BLA PNs, we expressed GCaMP6 specifically in PNs by injecting a CaMKII-dependent viral construct into the BLA of VIP-cre mice. Co-expression of cre-dependent ArchT-tdTomato allowed for simultaneous optogenetic manipulation of VIP BLA interneurons (**Figures 4H and S6A**). Similar to VIP interneurons, the first trials induced stronger US responses in BLA PNs of control animals expressing tdTomato in VIP cells (**Figure 4G, I**). In comparison, simultaneous inhibition of VIP interneurons with ArchT reduced PN US responses, as demonstrated by a significant decrease in US amplitude difference (light vs no-light trials) between control and ArchT mice (**Figure 4J-K**). These data show that VIP BLA interneuron activation can open a disinhibitory gate for neighboring PNs, leading to stronger somatic depolarization upon US application. Moreover, we analyzed the types of response patterns of BLA PNs during the predictive CS (Grewe et al., 2017; Gründemann et al., 2018). K-means clustering based on the Ca^2+^ activity during the last three CS trials of all n=545 cells from both control and ArchT groups revealed three distinct types of CS activity patterns in BLA PNs, with one group of CS responsive cells, which were found to show plastic responses when compared to the first three trials (**Figures 4L-M and S6C**). This fraction of CS-up cells was significantly smaller in the ArchT group, in which we suppressed VIP activity during the aversive US in the first three trials (**Figures 4N and S6C**), demonstrating that VIP US activity supports PN CS plasticity.

Finally, we addressed whether the observed US activation of VIP BLA interneurons is necessary for associative learning. We bilaterally expressed the inhibitory opsin ArchT specifically in VIP interneurons of the BLA and implanted optical fibers above the virus injection sites (**Figures 5A-B and S7B-D**). Mice were subjected to a single trial auditory fear conditioning paradigm, in which the activity of VIP interneurons was selectively suppressed during the aversive US using the same yellow light parameters as before (**Figures 5C and S7A**). Optogenetic inhibition of VIP BLA interneurons had no effect on freezing or locomotion of naïve mice (**Figure S7F-G**), nor did it affect the unconditioned response during the aversive US (**Figures 5D and S7H**). However, already during the acquisition phase, we observed significantly decreased freezing levels during the post-US period in mice with optogenetic inhibition of VIP activity during the US compared to control groups (**Figure 5E**). Most importantly, when testing fear memory after a 24 h consolidation phase, we found that optogenetic suppression of VIP interneuron US responses significantly prevented fear memory formation compared to controls (**Figure 5F**). Re-conditioning all groups of mice using a different auditory CS (CS2) without any optogenetic manipulation during the US (**Figure S7A, I-J**) revealed strong, CS2-specific freezing, confirming that the ArchT group was able to learn a CS-US association (**Figure 5G**). Together, these results demonstrate that sensory gating by activation of VIP BLA interneurons during the aversive US is necessary for associative learning.

**Figure 5.**
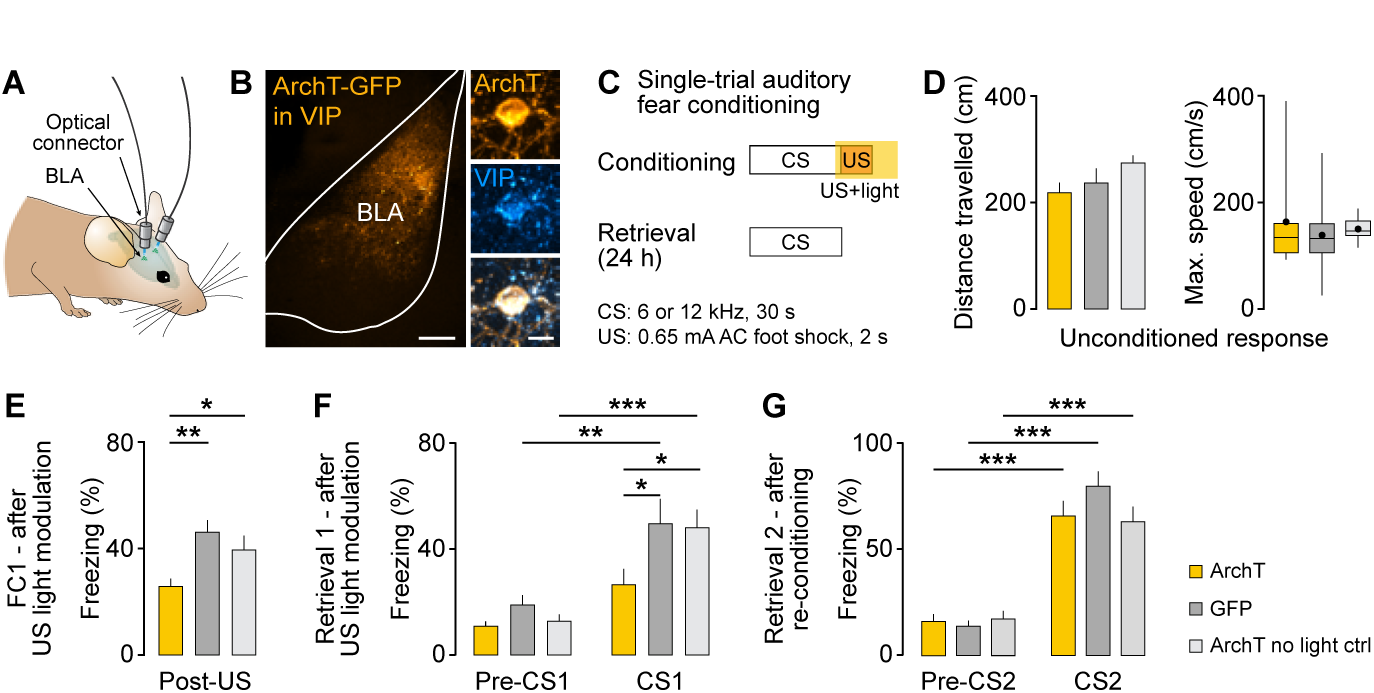
VIP BLA interneuron activation during the aversive US is necessary for learning. **(A)** Bilateral injection and fiber implantation scheme for optogenetic loss-of-function experiments in VIP-cre mice. **(B)** Expression of ArchT-GFP in the BLA of VIP-cre mice. High magnification image shows VIP immunolabelling in an ArchT-expressing cell. Scale bars, 200 μm and 20 μm (high magnification). **(C)** Schematic illustrating behavioral paradigm. VIP BLA interneuron activity was specifically suppressed during the US of the single-trial auditory fear conditioning paradigm. Re-conditioning was performed to a different CS2 without optogenetic manipulations during the US. **(D)** Neither distance travelled (left) nor maximum speed (right) during the US are affected by concomitant VIP BLA interneuron inhibition. Here and following: ArchT, N=14 mice; GFP, N=11; ArchT no light ctrl, N=12. **(E)** Post-US freezing is significantly diminished when VIP BLA interneurons during the US are optogenetically suppressed (one-way ANOVA, F=6.614, P<0.01; Holm-Sidak’s multiple comparisons test, ArchT vs GFP, P<0.01; ArchT vs ArchT no light ctrl, P<0.05). **(F)** Optogenetic inhibition of VIP BLA interneuron US activity during fear conditioning impairs associative learning as measured by freezing responses to CS1 on retrieval day (two-way ANOVA, main effect group, F_(2,68)_=1.859, P<0.05, main effect pre-CS1 to CS1, F_(1,68)_=36.47, P<0.001; Holm-Sidak’s multiple comparisons test, CS1 ArchT vs CS1 GFP, P<0.05; CS1 ArchT vs CS1 ArchT no light ctrl, P<0.05; Pre-CS1 vs CS1 GFP, P<0.01; Pre-CS1 vs CS1 ArchT no light ctrl, P<0.001). **(G)** Re-conditioning to a different CS2 without optogenetic manipulation of VIP BLA interneurons during the US does induce fear learning in all groups, but does not differ between ArchT and control mice (two-way ANOVA, main effect pre-CS2 to CS2, F_(1,68)_=136.9, P<0.001; Holm-Sidak’s multiple comparisons test, pre-CS2 vs CS2 ArchT, P<0.001; Pre-CS2 vs CS2 GFP, P<0.01; Pre-CS2 vs CS2 ArchT no light ctrl, P<0.001). Box-whisker plots show median values and 25^th^/75^th^ percentiles with 10^th^ to 90^th^ percentile whiskers, dots additionally indicate the mean. Other data is shown as mean and s.e.m. * P<0.05, ** P<0.01, *** P<0.001. Details of statistical analysis are listed in Table S3.

### VIP interneuron activity is modulated by expectation

Based on these findings and our previous observation that US responses of VIP interneurons decrease during conditioning (**Figure 1J**), we hypothesized that US-driven VIP interneuron activation may be modulated by expectation. Alternatively, the intra-session decrease of US responses might simply reflect a habituation-like process. To discriminate between these possibilities, we used a repeated fear conditioning paradigm in which mice experienced a second conditioning after a 24 h consolidation period (**Figure 6A-B**). Using our deep brain Ca^2+^ imaging approach, we were able to follow the activity of n=201 VIP BLA interneurons (from N=7 mice) across the two consecutive days (**Figures 6C-D and S7L**). Identical to our previous results, US responses decreased during the first day of training. However, they did not recover on the second day of training, but remained at a low level both in terms of response amplitudes and the fraction of responding cells (**Figure 6E-G**). Notably, after memory consolidation, VIP cells showed decreased US responses but stronger CS+ activation predictive of the aversive stimulus compared to the first training day (**Figure 6D-E**).

**Figure 6.**
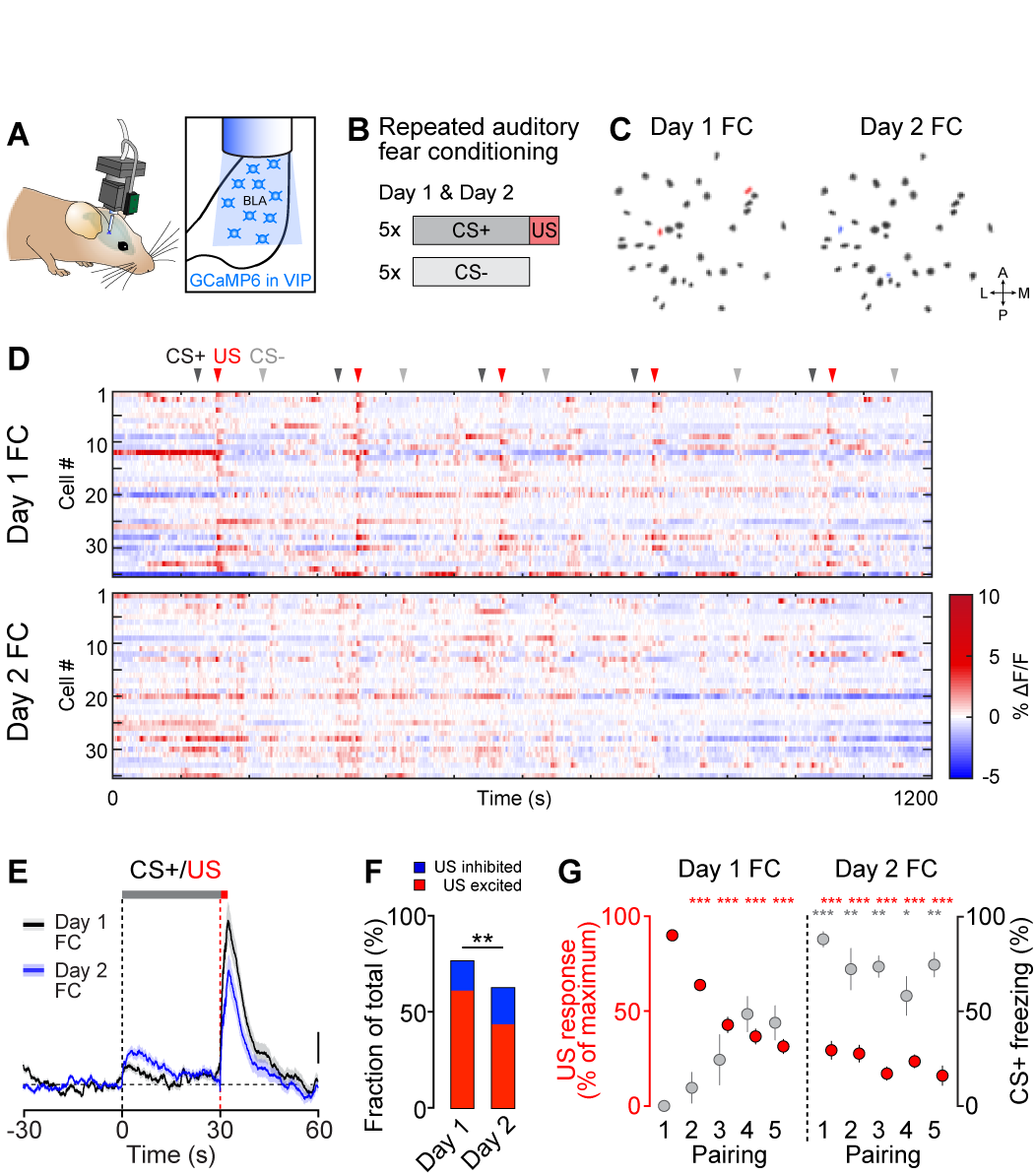
VIP BLA interneuron activity is modulated by expectation. **(A)** Schematic illustrating miniature microscope and implanted GRIN lens for deep brain Ca^2+^ imaging of BLA interneurons in freely behaving mice. **(B)** Schematic showing repeated auditory fear conditioning paradigm. CS+ and US pairings were presented alternating with the CS in an identical fashion on two consecutive days. **(C)** Spatial filters of identified VIP BLA interneurons (n=37 cells per day) on both recording days. Cells only found in the first fear conditioning session are labeled red (n=2), cells only found on the second day are blue (n=2). **(D)** Activity map of all VIP BLA interneurons identified on both consecutive days from the example mouse (n=35) during the two fear conditioning sessions. Arrowheads indicate onset of CS+ and US as well as intermingled CS, order of neurons is identical for both plots. Note that during day 1 and subsequently on day 2, cue-induced activity of VIP interneurons shifts from the aversive US to the predictive CS. **(E)** Average CS and US responses from all VIP BLA interneurons recorded during the repeated auditory fear conditioning paradigm (n=201 matched cells from N=7 mice, traces represent mean and s.e.m.). Scale bar, 0.2% ΔF/F. **(F)** The fraction of VIP BLA interneurons excited by the first aversive US decreases significantly on day 2 (Pearson’s X^2^ test, P0.01; Fisher’s exact test, US excited day 1 vs day 2, P<0.05; n=201 cells). **(G)** During day 1, US responses decrease over repeated pairings of CS+ and US in VIP BLA interneurons, while freezing during the CS+ increases in the same mice. After reconsolidation on day 2, when mice display a strong fear memory to the predictive CS+, US responses remain diminished (US response: Kruskal-Wallis test, H=322.4, P<0.0001; Dunn’s multiple comparisons test, D1 US1 vs D1 US2, P<0.0001 vs D1 US3, P<0.0001; vs D1 US4, P<0.0001; vs D1 US5, P<0.0001; vs D2 US1, P<0.0001; vs D2 US2, P<0.0001; vs D2 US3, P<0.0001; vs D2 US4, P<0.0001; vs D2 US5, P<0.0001; n=123 first US excited cells from N=7 mice; CS freezing: Kruskal-Wallis test, H=43.24, P<0.0001; Dunn’s multiple comparisons test, D1 CS1 vs D2 CS1, P<0.0001; vs D2 CS2, P<0.01; vs D2 CS3, P<0.01; vs D2 CS4, P<0.05; vs D2 CS5, P<0.01; N=7). Data in E and G is shown as mean and s.e.m. * P<0.05, ** P<0.01, *** P<0.001. Details of statistical analysis are listed in Table S3.

## DISCUSSION

Unexpected positive and negative experiences can be powerful triggers for associative memory formation. Although cortical VIP interneurons have been shown to be activated by salient stimuli (Pi et al., 2013), the causal involvement of inhibitory interneurons for associative learning instructed by these reinforcing cues remained so far unknown. In this study, using a combination of deep brain Ca^2+^ imaging and optogenetic manipulation of BLA interneurons during associative fear conditioning, we demonstrate for the first time that VIP interneurons are strongly activated by unexpected, aversive events during associative learning, and that this activation is a mandatory teaching signal for fear memory formation.

Within the local BLA microcircuit, we found that VIP BLA interneurons preferentially connect to other interneuron subtypes such as PN-targeting PV and SOM cells, which is in accordance with previous studies in BLA (Muller et al., 2003; Rhomberg et al., 2018) and cortical circuits (Lee et al., 2013; Pfeffer et al., 2013; Pi et al., 2013). Thus, during the aversive US presentation in associative fear learning when VIP interneurons are strongly activated, they are in an ideal position to release PNs from the powerful inhibition provided by SOM and PV interneurons. Remarkably, optogenetic inhibition of VIP interneuron activity during the aversive stimulus only mildly attenuated somatic US responses in PNs, but interfered significantly with the development of PN CS responses during learning (Grewe et al., 2017), a mechanism required for associative fear conditioning (Herry et al., 2008; Quirk et al., 1995). In line with this observation, our data support a model in which VIP BLA interneurons enable dendritic disinhibition in BLA PNs by mainly inhibiting SOM interneurons during aversive experiences. This effect is likely mediated in cooperation with the substantial fraction of US-excited PV BLA interneurons which also impinge on SOM interneurons, and is further promoted by the notion that PV and VIP BLA interneuron populations share similar long-range connectivity. In accordance with a dendritic disinhibition model and the observation that optogenetic inhibition of VIP interneuron activity during the US presentation only resulted in a mild decrease of somatic US responses in BLA PNs, the main effect on PN CS plasticity and ultimately learning could thus be driven independently of somatic activity by supporting dendritic plateau potentials in PNs (Gambino et al., 2014). Although necessary, the disinhibitory gating by VIP interneurons is likely not sufficient to induce learning, as it does not activate dendrites per se and consequently requires other signals such as a direct excitatory US input to PNs as well as neuromodulation (Johansen et al., 2014). On the other hand, precisely-timed perisomatic inhibition of PNs during the aversive US provided by PV BLA interneurons could further be beneficial for learning by supporting cellular mechanisms of spike-timing-dependent plasticity (Humeau et al., 2005; Shin et al., 2006), or might prevent fear generalization by constraining the memory engram (Morrison et al., 2016; Shaban et al., 2006).

In addition to the canonical VIP → PV and/or SOM → PN circuit motif, we observed reciprocal inhibitory connectivity between all three investigated BLA interneuron subtypes. Cortical interneuron populations with mutual inhibitory connections have been reported to change their activity patterns depending on external cues or locomotion, leading to differential responses in PN ensembles (Dipoppa et al., 2018; Kuchibhotla et al., 2017; Pakan et al., 2016). In the BLA, these circuit motifs could equivalently allow for flexible and dynamic routing of information modulated by different extrinsic inputs to the distinct interneuron subtypes as identified by our monosynaptic rabies tracings. This might be essential to ensure behavioral flexibility depending e.g. on the internal state of the animal, including physiological need states such as hunger or thirst (Burgess et al., 2016; Calhoon et al., 2018; Livneh et al., 2017), or external cues such as contextual signals (Kuchibhotla et al., 2017; Pakan et al., 2016).

While the source of the VIP US excitation remains so far unknown, we have identified several candidate brain regions for transmitting the information about the aversive cue. Using monosynaptic rabies tracings from VIP BLA interneurons, we found direct inputs from the dorsal midline and intralaminar thalamus as well as the insular cortex, which have all been implicated in transmitting foot shock information to the BLA (Lanuza et al., 2004; 2008; Sengupta and McNally, 2014). Furthermore, we observed a high number of rabies-labelled cholinergic cells in the basal forebrain, a neuronal population previously shown to be activated by aversive events (Hangya et al., 2015) and to drive the activity of cortical interneurons (Fu et al., 2014; Letzkus et al., 2011).

Intriguingly, VIP interneurons in cortical regions have been shown to be uniformly activated by both aversive and appetitive stimuli (Pi et al., 2013), and similar activity patterns have been observed e.g. in neuromodulatory systems such as dopaminergic or cholinergic populations (Bromberg-Martin et al., 2010; Lin and Nicolelis, 2008). While VIP BLA interneuron responses to rewarding cues remain so far unknown, BLA PNs can show similar activity patterns during aversive and appetitive conditioning (Belova et al., 2007; Shabel and Janak, 2009; Tye et al., 2008). Although recent work indicates that these opposite valences activate distinct populations of BLA PNs (Beyeler et al., 2016; Gore et al., 2015; Grewe et al., 2017; Kim et al., 2016), associative learning instructed by punishing or rewarding stimuli might share common mechanisms to induce neuronal plasticity. This implies that disinhibitory gating and/or neuromodulatory signaling would depend on salience irrespective of stimulus valence, while negative or positive valence could be assigned to distinct BLA PN populations by separate excitatory inputs.

Notably, we found that VIP interneuron responses during associative learning are highly plastic themselves, and shift from instructive to predictive stimuli upon memory formation. This is reminiscent of the changes in CS and US responses observed in other brain regions such as in midbrain dopamine neurons (Bromberg-Martin et al., 2010) and is thus consistent with the notion that VIP interneurons in the BLA reflect stimulus salience as a function of how well it is predicted. As a consequence, VIP interneuron activation forms part of a teaching signal gating the induction of neural plasticity and memory formation upon exposure to unexpected, but salient sensory cues. Further studies will be needed to address if this is an intrinsic plasticity mechanism in VIP BLA interneurons, or if this activation pattern is predominantly transmitted by external inputs, e.g. via a previously proposed multi-synaptic pathway from the periaqueductal grey (Johansen et al., 2010b; McNally et al., 2011).

Taken together, our findings identify a novel form of adaptive disinhibitory gating in a highly-specialized subgroup of inhibitory interneurons, which likely represents a key functional motif for associative learning in a dynamic, interconnected circuit. By detecting unexpected, meaningful environmental cues, VIP interneurons allow to filter important from irrelevant stimuli, and ensure appropriate behavioral adaptations to salient events by gating associative memory formation.

## METHODS

### Animals

All animal procedures were performed in accordance with institutional guidelines and were approved by the Veterinary Department of the Canton of Basel-Stadt and by the Austrian Animal Experimentation Ethics Board. VIP-ires-cre (Taniguchi et al., 2011), SOM-ires-cre (Taniguchi et al., 2011), PV-ires-cre (Hippenmeyer et al., 2005), PV-ires-flpE (Madisen et al., 2015) and SOM-ires-flpO (He et al., 2016) mice were used for cre or flp-dependent expression of viral vectors. Only heterozygous (cre/wt and flp/wt) mice were used for experiments, with the exception of cre or flp-negative littermates (wt/wt) for virus control injections. For some *in vitro* electrophysiology experiments, VIP-ires-cre mice were crossed with PV-ires-flpE or SOM-ires-flpO, as well as PV-ires-flpE with SOM-ires-cre lines, to generate VIP-ires-cre∷PV-ires-flpE, VIP-ires-cre∷SOM-ires-flpO and PV-ires-flpE∷SOM-ires-cre mice, respectively. For behavioral experiments, only male mice (aged 2-3 months at the time of injection) were used. Male and female mice (2-4 months at the time of injection) were utilized for rabies tracings, *in vitro* electrophysiology and immunohistochemistry. All lines were backcrossed to a C57BL/6J background for at least 7 generations. Mice were individually housed for at least two weeks before starting behavioral paradigms. Littermates of the same sex were randomly assigned to experimental groups. Animals were kept in a 12 h light/dark cycle with access to food and water *ad libitum.* All behavioral experiments were conducted during the light cycle.

### Surgical procedures and viral vectors

Mice were anesthetized using isoflurane (3-5% for induction, 1-2% for maintenance; Attane, Provet) in oxygen-enriched air (Oxymat 3, Weinmann) and fixed on a stereotactic frame (Model 1900, Kopf Instruments). Injections of buprenorphine (Temgesic, Indivior UK Limited; 0.1 mg/kg bodyweight subcutaneously 30 min prior to anesthesia) and ropivacain (Naropin, AstraZeneca; 0.1 ml locally under the scalp prior to incision) were provided for analgesia. Postoperative pain medication included buprenorphine (0.3 mg/ml in the drinking water; overnight) and injections of meloxicam (Metacam, Boehringer Ingelheim; 0.01 mg/kg subcutaneously) for up to three days if necessary. Ophthalmic ointment was applied to avoid eye drying. Body temperature of the experimental animal was maintained at 36°C using a feedback-controlled heating pad (FHC).

#### Deep brain imaging

AAV2/9.CAG.flex.GCaMP6s or AAV2/9.CAG.flex.GCaMP6f (400 nl, University of Pennsylvania Vector Core, UPenn) was unilaterally injected into the BLA of VIP-ires-cre, PV-ires-cre or SOM-ires-cre mice using a precision micropositioner (Model 2650, Kopf Instruments) and pulled glass pipettes (tip diameter about 20 um) connected to a Picospritzer III microinjection system (Parker Hannifin Corporation) at the following coordinates from bregma: AP −1.5 mm, ML 3.3 mm, DV 4.1-4.5 mm. For combined imaging and optogenetic manipulations of VIP interneurons, VIP-ires-cre mice were injected with AAV2/9.CAG.flex.GCaMP6s (400 nl) and AAV2/5.CAG.flex.ArchT-tdTomato (200 nl, University of North Carolina Vector Core, UNC). For combined imaging of BLA principal neurons and optogenetic manipulations of VIP interneurons, AAV2/5.CaMKII-GCaMP6f (400 nl, UPenn) was injected into the BLA with AAV2/5.CAG.flex.ArchT-tdTomato (200 nl, UNC) or AAV2/1.CAG.flex.tdTomato (200 nl, UPenn). Two weeks after virus injection, a gradient-index microendoscope (GRIN lens, 0.6 x 7.3 mm, GLP-0673, Inscopix) was implanted into the BLA as described previously (Xu et al., 2016). In brief, a sterile needle was used to make an incision above the imaging site. The GRIN lens was subsequently lowered into the brain with a micropositioner (coordinates from bregma: AP −1.6 mm, ML 3.2 mm, DV 4.5 mm) using a custom-build lens holder and fixed to the skull using UV light-curable glue (Henkel, Loctite 4305). Dental acrylic (Paladur, Heraeus) was used to seal the skull and attach a custom-made head bar for animal fixation during the miniature microscope mounting procedure. Mice were allowed to recover for one week after GRIN lens implantation before starting to check for GCaMP expression.

#### Optogenetic manipulations

VIP-ire-cre mice were bilaterally injected with AAV2/5.CAG.flex.ArchT-GFP (UNC) or AAV2/1.CAG.flex.GFP (UPenn) into the BLA (200 nl per hemisphere, coordinates from bregma: AP −1.5 mm, ML ±3.3 mm, DV 4.2-4.4 mm). Immediately after AAV injection, mice were bilaterally implanted with custom-made optic fiber connectors (fiber numerical aperture: 0.48, fiber inner core diameter: 200 um, Thorlabs). Fiber tips were lowered to −4 mm below cortical surface with a micropositioner. Implants were fixed to the skull with cyanoacrylate glue (Ultra Gel, Henkel) and miniature screws (P.A. Precision Screws). Dental acrylic mixed with black paint was used to seal the skull. Mice were allowed to recover for four weeks before behavioral training to ensure adequate virus expression.

#### Rabies tracings

VIP-ires-cre, PV-ires-cre or SOM-ires-cre mice were unilaterally injected with AAV2/7 or AAV2/1.Ef1a.DIO.TVA950-2A-CVS11G (50-100 nl, Vector Biolabs, custom production) into the BLA (coordinates from bregma: AP −1.5 mm, ML −3.3 mm, DV 4.4 mm). Data was pooled as no differences in viral efficiency were observed between the two AAV serotypes. To allow for sufficient expression of rabies glycoprotein and TVA receptor, mice were allowed to recover for two weeks before the injection of rabiesΔG-GFP-EnvA (RV-GFP) or rabiesΔG-RFP-EnvA (RV-RFP; 50-100 nl, custom production (Xu et al., 2016)).

#### In vitro electrophysiology

VIP-ires-cre∷PV-ires-flpE or VIP-ires-cre∷SOM-ires-flpO mice were bilaterally injected into the BLA with an AAV for ChR2 expression (200 nl per hemisphere, coordinates from bregma: AP −1.5 mm, ML ±3.3 mm, DV 4.2-4.4 mm). Two weeks later, an AAV for a fluorescent marker (400 nl) was bilaterally injected using the same coordinates. The following combinations of AAVs were used: For VIP→PV, VIP→SOM and SOM→PV: AAV2/5.EF1a.DIO.ChR2-EYFP (UPenn) with AAV2/5.hSyn.CRE-off-FLP-on.mCherry (Vector Biolabs, custom production). For PV→VIP, SOM→VIP and PV→SOM: AAVDJ.hSyn.CRE-off-FLP-on.ChR2-EYFP (UNC) with AAV2/1.CAG.flex.tdTomato (UPenn). Mice were allowed to recover for two more weeks before *in vitro* electrophysiology experiments. For functional tests of ArchT, AAV2/5.CAG.flex.ArchT-GFP or AAV2/5.CAG.flex.ArchT-tdTomato (UNC) was bilaterally injected into the BLA of VIP-ires-cre mice (200 nl) and animals were allowed to recover for four weeks.

### Immunohistochemistry

Mice were deeply anaesthetized with urethane (2 g/kg; intraperitoneally) and transcardially perfused with 0.9% NaCl followed by 4% paraformaldehyde in PBS. Brains were post-fixed for 2 h at 4°C and subsequently stored in PBS at 4°C. 80 **u**m coronal slices containing the BLA were cut with a vibratome (VT1000S, Leica). Sections were washed in PBS four times and blocked in 10% normal horse serum (NHS, Vector Laboratories) and 0.5% Triton X-100 (Sigma-Aldrich) in PBS for 2 h at room temperature. Slices were subsequently incubated in a combination of the following primary antibodies in carrier solution (1% NHS, 0.5% Triton X-100 in PBS) for 48 h at 4°C: rabbit anti-VIP (1:1000, Immunostar, 20077, LOT# 1339001), rat anti-SOM (1:500, Merck Millipore, MAB354, LOT# 232625), guinea pig anti-PV (1:500, Synaptic Systems, 195004, LOT# 195004/10), chicken anti-GFP (1:1000, Thermo Fisher Scientific, A10262, LOT# 1602788), chicken anti-RFP (1:500, Merck Millipore, AB3528, Lot# 2302143), mouse anti-CaMKII (1:500, Abcam, AB22609, Lot# GR220920-3), rabbit anti-2A peptide antibody (1:500, Merck Millipore, ABS31, LOT # 2746420) or mouse anti-2A peptide antibody (1:500, Novus Biologicals, NBP2-59627, LOT# A-1). After washing three times with 0.1 % Triton X-100 in PBS, sections were incubated for 12-24 h at 4°C with a combination of the following secondary antibodies (all 1:750 in carrier solution, Thermo Fisher Scientific): goat anti-chicken Alexa Fluor 488 (A11039, Lot# 1691381) or Alexa Fluor 568 (A11041, Lot# 1776042), goat anti-rabbit Alexa Fluor 488 (A11008, Lot# 1705869) or Alexa Fluor 647 (A21245, Lot# 1778005), goat anti-rat Alexa Fluor 568 (A11077, Lot# 692966) or Alexa Fluor 647 (A21247, Lot# 1524910), goat anti-guinea pig Alexa Fluor 647 (A21450, Lot# 1611324), goat-anti mouse Alexa Fluor 405 (A41553, Lot# 1512096) or Alexa Fluor 647 (A21235, Lot# 1890864). After washing four times in PBS, sections were mounted on glass slides and coverslipped. Sections were scanned using a laser scanning confocal microscope (LSM700, Carl Zeiss AG) equipped with a 10x air objective (Plan-Apochromat 10x/0.45) or 20x air objective (Plan-Apochromat 20x/0.8). Tiled z-stacks (3 um) of the BLA were acquired and stitched with Zeiss software processing tool (ZEN 2.3, black edition, Carl Zeiss AG). Images were imported in Imaris Software (Bitplane AG) to count somata expressing the fluorophore only or co-expressing the fluorophore and the peptide/protein of interest. Every third section containing the BLA was analyzed.

### Deep brain calcium imaging

#### Miniature microscope imaging

Two to four weeks after GRIN lens implantation, mice were head-fixed to check for sufficient expression of GCaMP6 using a miniature microscope (nVista HD, Inscopix). Mice were briefly anesthetized with isoflurane to fix the microscope baseplate (BLP-2, Inscopix) to the skull using light curable glue (Vertise Flow, Kerr). The microscope was removed and the baseplate was capped with a baseplate cover (Inscopix) whenever the animal was returned to its home cage. The microscope was mounted on a daily basis immediately before starting the behavioral session. Mice were habituated to the brief head-fixation on a running wheel for miniature microscope mounting for at least three days before the behavioral paradigm. Imaging data was acquired using nVista HD software (Inscopix) at a frame rate of 20 Hz with an LED power of 40-80% (0.9-1.7 mW at the objective, 475 nm), analogue gain of 1-2 and a field of view of 650 x 650 μm. For combined imaging and optogenetic experiments, the nVoke imaging system and software (Inscopix) were used. Imaging LED power was set to 0.5-0.8 (0.4-0.7 mW at the objective, 448 nm) with an analogue gain of 1.5-3. For individual mice, the same imaging parameters were kept across repeated behavioral sessions. Timestamps of imaging frames and behavioral stimuli were collected for alignment using the MAP data acquisition system (Plexon).

#### Discriminative fear conditioning paradigm

Two different contexts were used for the associative fear learning paradigm. Context A (retrieval context) consisted of a clear cylindrical chamber (diameter: 23 cm) with a smooth floor, placed into a dark-walled sound attenuating chamber under dim light conditions. The chamber was cleaned with 1% acetic acid. Context B (fear conditioning context) contained a clear square chamber (26.1 x 26.1 cm) with an electrical grid floor (Coulbourn Instruments) for foot shock delivery, placed into a light-colored sound attenuating chamber with bright light conditions, and was cleaned with 70% ethanol. Both chambers contained overhead speakers for delivery of auditory stimuli, which were generated using a System 3 RP2.1 real time processor and SA1 stereo amplifier with RPvdsEx Software (all Tucker-Davis Technologies). A precision animal shocker (H13-15, Coulbourn Instruments) was used for the delivery of alternating current (AC) foot shocks through the grid floor. Behavioral protocols for stimulus control were generated with Radiant Software (Plexon) via TTL pulses. On day 1, mice were habituated in context A. Two different pure tones (conditioned stimulus, CS; 6 kHz and 12 kHz, total duration of 30 s, consisting of 200 ms pips repeated at 0.9 Hz; 75 dB sound pressure level) were presented five times each in an alternated fashion with a pseudorandom ITI (range 30-90 s, 2 min baseline before first CS). On day 2, mice were conditioned in context B to one of the pure tones (CS+) by pairing it with an unconditioned stimulus (US; 2 s foot shock, 0.65 mA AC; applied after the CS at the time of next expected pip occurrence). The other pure tone was used as a CS and not paired with a US. CS+ with US and CS were presented alternating five times each in a pseudorandom fashion (ITI 60-90 s), starting with the CS+ after a 2 min baseline period. Animals remained in the context for 1 min after the last CS-presentation and were then returned to their home cage. On day 3, fear memory was tested in context A. After a 2 min baseline period, the CS was presented four times, followed by 12 CS+ presentations (ITI 60-90 s). A second group of VIP-cre mice was re-exposed to the fear conditioning paradigm on day 3 and tested for fear memory in the retrieval context on day 4 as described above. The use of 6 kHz and 12 kHz as CS+ was counterbalanced in individual groups.

#### Fear conditioning for combined deep brain imaging and optogenetic manipulations

Two different contexts were used as described above. On day 1, mice were habituated in context A. A 6 kHz CS (total duration of 30 s, 200 ms pips repeated at 0.9 Hz; 75 dB sound pressure level) was presented six times with a pseudorandom ITI (range 30-90 s, 2 min baseline before first CS). Subsequently, three yellow light stimuli were applied using the nVoke system (4.5 s continuous illumination, 590 nm LED, 12 mW at the objective of the miniature microscope, ITI 30 s). On day 2, mice were conditioned to the CS (2 min baseline) by pairing it with a US (2 s foot shock, 0.65 mA AC; applied after the CS at the time of next expected pip occurrence). Six repetitions of CS and US pairings were applied. The first three US presentations were combined with yellow light (4.5 s, starting 500 ms before the US onset), while the last three US presentations were used as no-light controls. Animals remained in the context for 1 min after the last US and were then returned to their home cage. On day 3, fear memory was tested in context A where mice were exposed to 12 CS presentations (ITI 60-90 s) after a 2 min baseline period. Animals further received yellow light stimulations after the CS presentations (4.5 s, 10 s, 20 s, four times each, ITI 30-60 s).

#### Verification of implant sites

Upon completion of the behavioral paradigm, mice were transcardially perfused (as above). The GRIN lens was removed and brains post-fixed in 4% paraformaldehyde for at least 2 h at 4°C. Coronal sections (120 nm) containing the BLA were cut with a vibratome (VT1000S), immediately mounted on glass slides and coverslipped. To verify the microendoscope position, sections were scanned with a laser scanning confocal microscope (LSM700) equipped with a 10x air objective (Plan-Apochromat 10x/0.45) and matched against a mouse brain atlas (Paxinos and Franklin, 2001). Mice were post-hoc excluded from the analysis if the GRIN lens was placed outside of the BLA. Forn Voke experiments, mice were further excluded if they did not exhibit BLA-specific ArchT or tdTomato expression.

#### Analysis of behavior

All behavioral sessions were recorded using an overhead camera and Cineplex software (Plexon). Mice were tracked using contour tracking, and freezing behavior was automatically scored with the assistance of a frame-by-frame analysis of pixel change (Cineplex Editor, Plexon). Freezing behavior minimum duration threshold was set to 2 s. Automatically detected freezing was cross-checked on the video recording to eliminate false-positive and false-negative freezing bouts (e.g., during grooming episodes or due to motion artefacts caused by cable movement, respectively). To extract the animals’ speed (cm/s) during the behavioral paradigm, the size of the behavioral chamber was calibrated with the pixel dimension of the camera. Freezing and speed data were processed with custom-written Matlab (Mathworks) scripts.

#### Analysis of calcium imaging data

Imaging frames were normalized across the whole frame by dividing each frame by a Fast Fourier Transform band pass-filtered version of the frame using ImageJ (NIH) (Schneider et al., 2012). XY movement was corrected using the TurboReg ImageJ plugin (Thevenaz et al., 1998). Further analysis was conducted using Matlab. Spatial filters for individual neurons were defined using an automated cell sorting routine based on the entire imaging session using principal and independent component analyses (Mukamel et al., 2009). Every cell included in the analyses was confirmed by visual inspection and spatial filters that did not correspond to neurons (e.g., blood vessels) were discarded. Cell masks were then applied to the movie to obtain raw calcium fluorescence. Relative changes in calcium fluorescence F were calculated by ΔF/F0 = (F - F0)/F0 (with F0 = mean fluorescence of entire trace). For repeated fear conditioning experiments, the cell map obtained from day 2 was aligned to the day 1 reference map using TurboReg. Stimulus responses were baselined to the 30 s pre-event period for figure display. To define responsive cells, average time-binned Ca^2+^ signals were compared between the stimulus and equivalent baseline period using a Wilcoxon rank sum test with a significance threshold of *P*<0.05. Stimulus response period was set to 4 s for US (8 time bins of 0.5 s) and 30 s for CS presentations (30 time bins of 1 s). To characterize PN response dynamics to CS presentation during conditioning upon VIP interneuron US modulation, we collected responses in a time window starting 10 s before stimulus CS and ending with CS offset and averaged across the last three presentations of the CS (non-manipulated US). K-means clustering was performed using all 545 cells from both control and VIP-ArchT groups with the cosine distance function (k=5), and clusters were manually characterized as the 3 response types (1) CS responsive cells, which were found to show plastic responses when compared to the first three trials, as well as CS non-responsive cells with either (2) activity during both baseline and CS or (3) no activity during baseline and CS (**Figure S6C**).

### Optogenetic manipulation of behavior

Optogenetic manipulation experiments and analysis of freezing were performed by an experimenter blinded to the group assignment of the animal (both virus condition and light condition). Animals were allocated to experimental groups without pre-determined criteria and could be later identified by unique markers for group assignment. Before behavioral experiments, all mice were habituated to the experimenter and to the optical fiber connection procedure by handling and short head-restraining for at least three days. On the experimental days, implanted fibers were connected to a custom-built laser bench (Life Imaging Services) with custom fiber patch cables. An acusto-optic tunable filter (AOTFnC-400.650-TN, AA Opto-Electronic) controlled laser intensity (MGL-F-589, 589 nm wavelength, CNI lasers). At the beginning of the behavioral session, laser power at the tip of the fiber patch cables was tested with an optical power and energy meter (PM100D, ThorLabs) and adjusted to an intensity of 15 mW at the fiber patch cable tips, equating approximately 12 mW at the implanted fiber ends.

Mice were subjected to a single-trial auditory fear conditioning paradigm (**Figure S7A**). Two different contexts were used as described above. On day 1, mice were habituated in context A (retrieval context). Following a baseline period (4 min), two different CSs (CS1 and CS2; pip frequency: 12 kHz and 6 kHz, total CS duration of 30 s, consisting of 200 ms pips repeated at 0.9 Hz; 75 dB sound pressure level) were presented four times in an alternated fashion with a pseudorandom ITI (range 30-90 s). After the last CS, animals received four presentations of a yellow light stimulus (4.5 s continuous illumination, ITI 30 s). On day 2 (fear conditioning, context B), following a baseline period (4 min), animals were exposed to one CS (CS1: either 12 kHz or 6 kHz) paired with a US (2 s foot shock, 0.65 mA AC; applied after the CS at the time of next expected pip occurrence). The US was paired with yellow light stimulation (4.5 s, starting 500 ms before US onset). Mice remained in the context for 1 min after the US presentation and were then returned to their home cage. On day 3 (retrieval 1), fear memory was tested in context A by presenting the CS1 (4 min baseline) without any light stimulation or reinforcement. Freezing behavior induced by the presentation of CS1 was used to determine the effect of the optogenetic manipulation on fear learning in comparison to control groups. As an additional control, on day 4 (re-conditioning) mice were placed back to context B where they were conditioned by paring the CS2 with the US in absence of any light stimulation (same as above, using CS2: 6 kHz or 12 kHz, depending on CS1 on day 2). Fear memory to CS2 was tested on day 5 in context A (4 min baseline). After CS presentation, animals received four 4.5 s and two 10 s presentations of the light stimulus (ITI 20 s). The order of 12 kHz and 6 kHz for both conditioning paradigms was counterbalanced within behavioral groups. For the “ArchT no light” control group, the paradigm was unaltered except for the removal of any light delivery on day 1, day 2 and day 5.

Behavior was analyzed as described above by an observer blind to the group assignment (both virus condition and light condition). Freezing responses to yellow light during habituation (day 1) and after retrieval 2 (day 5) were quantified during (4.5 s or 10 s) and after (30 s inter-stimulus interval) stimulus presentation. Running speed in naïve mice was measured before (10 s pre-laser interval), during (4.5 s stimulation), and after (10 s interval post-laser) four repeated yellow light stimulations during the habituation session. Post-shock freezing was assessed for a 30 s period after US delivery. Freezing during the CS period (30 s) was compared to the pre-CS period (4 min baseline) to evaluate learning.

#### Verification of injection sites and optical fiber placement

Upon completion of the behavioral paradigm, mice were transcardially perfused (as above) and optical fibers removed. Brains were post-fixed in 4% paraformaldehyde for at least 2 h at 4 °C and cut in 80|xm coronal slices with a vibratome (VT1000S). Sections containing the BLA were immediately mounted on glass slides and coverslipped. To verify specificity of viral expression and fiber tip placement, sections were scanned with a 10x air objective (Plan-Apochromat 10x/0.45) using a laser scanning confocal microscope (LSM700). Fiber tip placements were matched against a mouse brain atlas (Paxinos and Franklin, 2001). Animals were post-hoc excluded from the analysis if (1) US delivery failed during any of the fear conditioning sessions (day 2 or day 4), (2) they did not show bilateral expression of the virus, (3) major virus expression was detected outside of the BLA or (4) they did not exhibit correct fiber placement (no more than 300 μm away from the BLA, **Figure S7D**).

### Monosynaptic rabies tracing

#### Immunohistochemistry

One week after rabies virus injection, mice were transcardially perfused (as above). After post-fixation in 4% paraformaldehyde for 2 h at 4°C, brains were embedded in 4% agarose in PBS and cut into 80 **|x**m coronal sections from rostral to caudal (+4.28 to −7.08 AP from bregma) with a vibratome (VT1000S). After washing three times with PBS, every third section from each brain was incubated in blocking solution (3% bovine serum albumin (Sigma-Aldrich) in 0.5% Triton X-100 in PBS) for 2 h at room temperature. Subsequently, sections were incubated with primary rabbit anti-2A peptide antibody (1:500, Merck Millipore, ABS31, LOT # 2746420) in carrier solution (as above) for 48 h at 4 °C. Samples were rinsed with 0.1 % Triton X-100 in PBS three times and then incubated overnight at 4°C with donkey anti-rabbit Alexa Fluor 568 (1:750, Thermo Fisher Scientific, A10042, LOT# 1757124) or donkey anti-rabbit Alexa Fluor 647 (1:750, Thermo Fisher Scientific, A31573, LOT# 1786284) in carrier solution. Finally, sections were washed four times with PBS, mounted on glass slides and coverslipped. The remaining sections were either directly mounted on slides and coverslipped, or processed for further immunohistochemical analysis. In some brains (N=3 from VIP-cre), one third of cut slices were incubated, as described above, first in blocking solution (10% normal horse serum (NHS) and 0.5% Triton X-100 in PBS) and then with goat anti-ChAT antibody (1:250, Merck Millipore, AB144P, LOT# 2147041) in 1% NHS and 0.5% Triton X-100 in PBS, followed by donkey anti-goat Alexa Fluor 647 secondary antibody (1:750, Thermo Fisher Scientific, A21447, LOT# 1841382).

#### Image acquisition

All sections from each brain were imaged using an Axioscan Z1 slide scanner (with ZEN 2.3, blue edition, Carl Zeiss AG) with a 5x air objective (Fluar 5x/0.25). Selected brain slices, including sections containing the BLA, were further imaged using a laser scanning confocal microscope (LSM700) with 10x air objective (Plan-Apochromat 10x/0.45). Tiled z-stacks (3 um) of the region of interest were acquired and stitched with Zeiss software processing tool (ZEN 2.3, black edition). Tiled images of the BLA were imported into Imaris Software to analyze co-expression profiles of 2A peptide. In brief, sections were inspected through the whole z-stack and 2A peptide/RV double-positive somata were marked with Imaris spot function and automatically counted as spots. A cell was considered a starter cell if it co-expressed 2A peptide and RV. Total starter cell number in the entire BLA was extrapolated by counting double-positive 2A peptide/RV cells in 1/3 of BLA sections. Brains were excluded from the analysis if more than 10% of starter cells were found outside the BLA.

#### Data analysis

Images from individual brain sections were sorted in the correct order from rostral to caudal and aligned with the TrakEM plug-in for ImageJ to create a serial z-stack. Regions of interest (ROI) were manually drawn in ImageJ using a mouse brain atlas (Paxinos and Franklin, 2001) for reference and saved as a mask with a custom written ImageJ macro (Facility for Advanced Imaging & Microscopy, FMI). A prediction model for automatic somata detection was created with Ilastik software (Version 1.1.5) for each analyzed brain. A custom Matlab script (Facility for Advanced Imaging & Microscopy, FMI) based on z-stack, ROI masks and prediction model of each brain was used to automatically quantify RV^+^ cells in each brain area. Visual inspection of Matlab script images output was used to correct for false positive and false negative cells. The fraction of inputs for each brain area was calculated as the percentage of presynaptic cells per ROI over the sum of the absolute number of presynaptic cells in all counted brain regions. The convergence index (CI) for each brain area was calculated as the ratio between the number of detected RV^+^ cells per brain area and the number of starter cells in the BLA. For quantification within selected subregions, only the input neurons ipsilateral to the injection site were taken into consideration. Areas that contained less than 1 % of the total sum inputs and the BLA itself were excluded. Brain areas under the following denominations included specific subareas (anatomical abbreviations used for figure display): (1) insular cortex (Ai): agranular insular cortex, dorsal (AiD) and ventral part (AiV); (2) basal forebrain (BF): ventral pallidum (VP), nucleus of the horizontal limb of the diagonal band (HDB), magnocellular preoptic nucleus (MCPO), substantia innominata (SI), basal nucleus of Meynart; (3) rhinal cortices (RhC): ectorhinal and perirhinal cortex, dorsal and ventral intermedial, dorsolateral entorhinal cortex (4) auditory cortex (AuC): primary auditory cortex (Au1), secondary auditory cortex, dorsal (AuD) and ventral part (AuV), temporal association cortex (TeA); (5) auditory thalamus (AuT): medial geniculate nucleus (medial part; MGM), suprageniculate thalamic nucleus (SG); (6) dorsal midline thalamus (dMT): paraventricular thalamic nucleus (PVT), paratenial nucleus (PT). Further presynaptic brain areas (anatomical abbreviations) included in the analysis were the ventral hippocampus (vHC), piriform cortex (Pir), cortex-amygdala transition zone with posterior cortical amygdaloid area (CxA), ventromedial hypothalamus (VMH), nucleus of the lateral olfactory tract (LOT), posterior intralaminar thalamus (PIL), dorsal raphe nucleus (DR) and medial orbital cortex (MO). Hierarchical cluster analysis of cases (injected animals per interneuron subtype) was based on fraction of inputs of each identified presynaptic brain area. Cluster method between groups linkage was based on measured squared Euclidean distance. Animals were classified according to the percentage of starter cell population detected in the lateral amygdala (LA). The cut-off percentage used for identifying preferential LA tracing injections was set to ≥70% LA starter cells, for preferential basal amygdala (BA) injections threshold was set to ≤30% LA starter cells. Other animals were considered as mixed LA/BA tracings.

### *In vitro* electrophysiology

#### Connectivity assays

Mice were deeply anaesthetized (ketamine 250 mg/kg and medetomidine 2.5 mg/kg bodyweight intraperitoneal) and transcardially perfused with ice-cold slicing ACSF (in mM: 124 NaCl, 2.7 KCl, 26 NaHCO_3_, 1.25 NaH_2_PO_4_, 2.5 glucose, 50 sucrose, 0.1 CaCl_2_, 6 MgCl_2_, 3 kynurenic acid, oxygenated with 95% O_2_/5% CO_2_, pH 7.4). The brain was rapidly removed from the skull, and coronal brain slices (300 μm) containing the BLA were prepared in ice-cold slicing ACSF with a vibrating-blade microtome (HM650V, Microm) equipped with a sapphire blade (Delaware Diamond Knives). For recovery, slices were kept in the dark for 45 min at 37°C in an interface chamber containing recording ACSF (in mM: 124 NaCl, 2.7 KCl, 26 NaHCO_3_, 1.25 NaH_2_PO_4_, 18.6 glucose, 2.25 ascorbic acid, 2 CaCl_2_, 1.3 MgCl_2_, oxygenated with 95% O_2_/5% CO_2_, pH 7.4) and afterwards at room temperature (20-22°C) until start of recordings. Experiments were performed in a submerged chamber on an upright microscope (BX50WI, Olympus) superfused with recording ACSF (as above, except: 2.5 mM CaCl_2_, 10 μM CNQX and 10 μM CPP) at a perfusion rate of 2-4 ml/min at 32°C. EYFP^+^/tdTomato^+^/mCherry^+^ interneurons were visualized using epifluorescence and a 40x water immersion objective (LumPlanFl 40x/0.8, Olympus).

Patch electrodes (3-5 MΩ) were pulled from borosilicate glass tubing and filled with internal solution (for voltage-clamp recordings in mM: 110 CsCl, 30 K-gluconate, 1.1 EGTA, 10 HEPES, 0.1 CaCl_2_, 4 Mg-ATP, 0.3 Na-GTP, 4 QX-314 chloride and 0.4% biocytin, pH 7.3; for current-clamp recordings in mM: 130 K-methylsulfate, 10 HEPES, 10 Na-phosphocreatine, 4 Mg-ATP, 0.3 Na-GTP, 5 KCl, 0.6 EGTA and 0.4% biocytin, pH 7.3). Voltage-clamp recordings were acquired in whole-cell mode at a holding potential of −70 mV. Conductance was determined with additional holding potentials of −60 mV and −50 mV. ChR2 expressing interneurons were photostimulated using a blue LED (PlexBright Blue, 465 nm, with LED-driver LD-1, Plexon) connected to an optical fiber positioned above the BLA. Five blue light pulses of 10 mW with 10 ms duration were applied at a frequency of 1 Hz. Inhibitory postsynaptic currents were averaged across at least 20 light pulses. In some slices, picrotoxin (100μM) was administered with the recording ACSF for the last recorded cell. In current-clamp recordings, spikes were evoked from a holding potential of about −60 mV with current injections for 500 ms, and the same current step was subsequently paired with blue light stimulation to activate ChR2-expressing interneurons. Spike probability of individual neurons with and without blue light stimulation was calculated from ten repeated trials. Functionality of ChR2 constructs used for slice electrophysiology was tested in on-cell and whole-cell mode (10 ms and 300 ms blue light pulses). Data was acquired with a Multiclamp 700A amplifier, Digidata 1440A A/D converter and pClamp 10 software (all Molecular Devices) at 20 kHz and filtered at 4 kHz (voltage-clamp) or 10 kHz (current-clamp). Whole-cell recordings were excluded if the access resistance exceeded 25 MΩ or changed more than 20% during the recordings. Data was analyzed using IGOR Pro software (Version 6.35A5, WaveMetrics) with NeuroMatic plug-in (Rothman and Silver, 2018). All chemicals were purchased from Sigma-Aldrich except for CNQX, CPP and QX-314 (Tocris Bioscience).

#### ArchT light sensitivity

To confirm functionality of the ArchT-GFP and ArchT-tdTomato constructs used for optogenetic manipulation experiments, whole-cell current-clamp recordings (as above) from GFP^+^/tdTomato^+^ VIP interneurons were performed in recording ACSF (as above, except 10 μM CNQX, 10μM CPP and 100μM picrotoxin). An optical fiber connected to a yellow LED (PlexBright Yellow, 590 nm, 2.5 mW maximum output, with LED-driver LD-1, Plexon) was positioned above the BLA. ArchT functionality was tested with 500 ms and 4.5 s yellow light pulses. Further, depolarizing current steps of 500 ms duration were applied from a holding potential of −60 mV (25 pA steps starting from −100 pA). The same current steps were subsequently paired with yellow light to assess the effect of ArchT activation on spiking. An identical protocol was used to test the effect of blue light (473 nm, 0.8 mW) on ArchT activation. Data from ArchT-GFP and ArchT-tdTomato constructs was pooled for analysis of ArchT light sensitivity.

#### Immunohistochemistry of patch slices

All cells were filled with biocytin during patch-clamp recordings. Outside-out patches were pulled at the end of each recording and slices were fixed for 1 h in 4% paraformaldehyde in PBS for subsequent immunohistochemistry. Slices were stored in PBS overnight (4°C). Free-floating sections were then washed with PBS three times before incubation in blocking solution (3% normal goat serum, 1% bovine serum albumin and 0.5% Triton X-100 in PBS) for 3 h at room temperature. Slices were then incubated for 48 h at 4°C in carrier solution (same as above) with a combination of the following primary antibodies: rabbit anti-VIP (1:1000, Immunostar, 20077, LOT# 1339001), rat anti-SOM(1:500, Merck Millipore, MAB354, LOT# 232625), guinea pig anti-PV (1:500, Synaptic Systems, 195004, LOT# 195004/10), chicken anti-GFP (1:1000, Thermo Fisher Scientific, A10262, LOT# 1602788). Sections were washed with 0.1 % Triton X-100 in PBS three times before adding secondary antibodies in carrier solution (all 1:750) for 24 h at 4°C: goat anti-chicken Alexa Fluor 488 (A11039, Lot# 1691381), goat anti-rabbit Alexa Fluor 647 (A21245, Lot# 1778005), goat anti-rat Alexa Fluor 647 (A21247, Lot# 1524910), goat anti-guinea pig Alexa Fluor 647 (A21450, Lot# 1611324; all Thermo Fisher Scientific) and streptavidin conjugated to Alexa Fluor 405 (1:1000, Thermo Fisher Scientific, S32351, Lot# 1712187). Finally, slices were washed in PBS four times, mounted and coverslipped. Z-stacks (1.5 um) of biocytin-filled cells were acquired using a confocal microscope (LSM700) with 63x objective (Plan-Apochromat 63x/1.40 Oil DIC) and 2-fold digital zoom. Cells were excluded from the analysis if (1) they were not recovered with immunohistochemistry or (2) if post-hoc immunohistochemistry did not confirm PV or SOM expression for recorded BLA interneurons or spiny dendrite morphology and large cell size for BLA principal cells.

### Statistical analysis and data presentation

All datasets were tested for Gaussian distribution using a Shapiro-Wilk normality test. Two datasets were statistically compared using a Student’s t test and the data values are expressed as mean and s.e.m. if the null hypothesis of normal distribution was not rejected. A two-way ANOVA was used when comparing more than two normally-distributed datasets. Post-hoc multiple comparisons were performed using the Holm-Sidak correction. If the null hypothesis of normal distribution was rejected, two datasets were compared using a Mann-Whitney U test and are presented as median values and 25^th^/75^th^ percentiles. Figures additionally display 10^th^ to 90^th^ percentiles. Pairwise comparisons were calculated with a Wilcoxon matched-pairs signed-rank test. Nonparametric comparison of datasets with more than two groups was carried out with a Kruskal-Wallis test and Dunn’s post-hoc correction. Connectivity ratios were compared using Pearson’s X^2^ test. In case of a significant result, a Fisher’s exact test was calculated between individual groups. Statistical analysis was carried out using Matlab or Prism 7 (GraphPad Software). Hierarchical cluster analysis for rabies virus tracing was carried out using IBM SPSS statistics version 24. Statistical significance threshold was set at 0.05 and significance levels are presented as * (*P*<0.05), ** (*P*<0.01) or *** (*P*<0.001) in all figures. Statistical tests and results are reported in the respective figure legends and Table S3. The number of analyzed cells is indicated with ‘n’, while ‘N’ declares the number of animals. No statistical methods were used to predetermine sample sizes. The sample sizes were chosen based on published studies in the field. Optogenetic manipulation experiments and analysis of behavior were performed by an experimenter blinded to the group assignment of the animal (both virus condition and light condition). Contrast and brightness of representative example images were minimally adjusted using ImageJ. For figure display, confocal images were further scaled (0.5×0.5) and electrophysiological traces were resampled to 5 kHz.

## ACKNOWLEDGEMENTS

We thank all members of the Lüthi and Ferraguti labs for helpful discussions and comments. We would like to thank P. Argast, P. Buchmann, T. Eichlisberger, A. Kovacevic, T. Lu, C. Müller and all staff of the FMI Animal Facility for excellent technical assistance. We further thank the Facility for Imaging and Microscopy at the FMI, in particular S. Bourke and R. Thierry, and the FMI IT department, in particular D. Flanders, S. Grzybek and R. Milani, for their support with data acquisition and analysis, as well as M. Stadler for statistical advice. We are grateful to the GENIE Program at Janelia Research Campus of the Howard Hughes Medical Institute for making GCaMP6 material available, C. Ramakrishnan and K. Deisseroth for viral constructs, and S. Arber and Z. J. Huang for sharing mouse lines. We further thank Inscopix for providing access to the nVoke integrated imaging and optogenetics system. This work was supported by the European Research Council (ERC) under the European Union’s Horizon 2020 research and innovation programme (grant agreement No 669582), by the National Center of Competences in Research: “SYNAPSY - The Synaptic Bases of Mental Diseases” (financed by the Swiss National Science Foundation, SNSF) and an SNSF core grant (all to AL); by the Austrian Science Fund (Fonds zur Förderung der Wissenschaftlichen Forschung) Sonderforschungsbereich grant F44-17-B23 and W012060-10 (to FF); as well as by a Young Investigator Grant from the Brain & Behavior Research Foundation (to SK); a NENS exchange grant (to EP); EMBO Long-Term Fellowships (to CX and JG); an SNSF Ambizione grant and SNSF Professorship (to JG); and by the Novartis Research Foundation.

## AUTHOR CONTRIBUTIONS

SK, EP, JG, FF and AL designed the project. SK, JG and YB performed and analyzed deep brain imaging experiments. EP and SD performed and analyzed rabies tracings. SK performed and analyzed *in vitro* electrophysiology. EP performed and analyzed optogenetic manipulation experiments. SK, EP and SD performed and analyzed immunohistochemistry. CX, KY and MM designed, generated and validated viral constructs. SK, EP, JG, FF and AL wrote the manuscript. All authors contributed to the experimental design, interpretation of the data and commented on the manuscript.

## Supplementary material

Adaptive disinhibitory gating by VIP interneurons permits associative learning Krabbe, Paradiso et al. 2018

**Figure S1.**
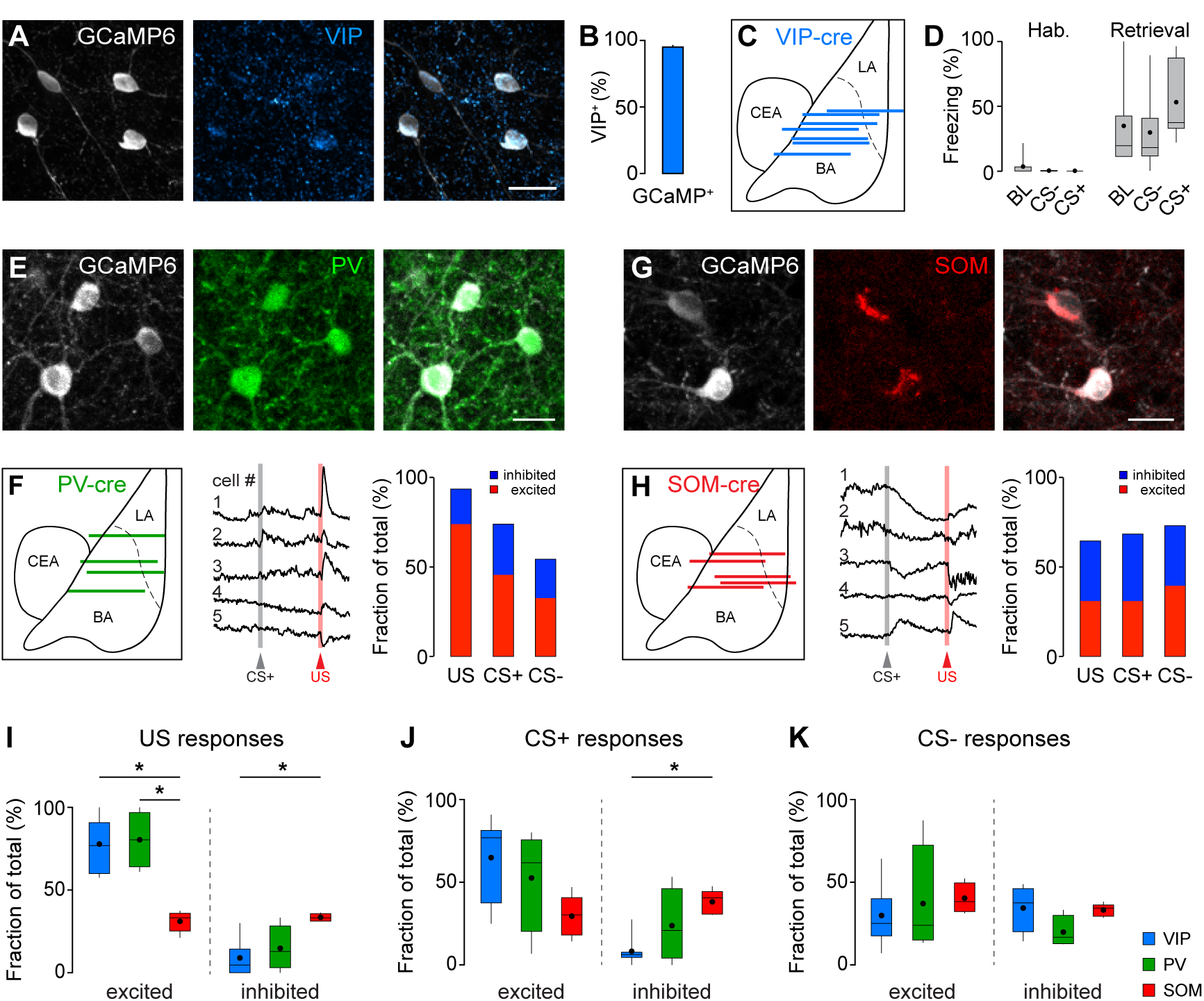
Heterogeneity of CS and US responses in BLA interneuron subtypes. **(A)** Representative example image of GCaMP6 expression in VIP interneurons in the BLA of VIP-cre mice. Scale bar, 20 μm. **(B)** Quantification of co-localization of viral GCaMP6 expression with VIP detected by immunohistochemistry (N=3 mice). **(C)** Schematic illustrating reconstructed implant sites of GRIN lenses (blue lines) within the BLA of VIP-cre mice for deep brain imaging experiments presented in Figure 1 matched to a mouse brain atlas (N=7 mice). LA, lateral amygdala; BA, basal amygdala; CEA, central amygdala. **(D)** Freezing levels before and after fear conditioning in GRIN lens-implanted VIP-cre mice (N=7). **(E)** Representative example image of GCaMP6 expression in PV interneurons in the BLA of PV-cre mice. Scale bar, 20 μm. **(F)** Left to right, GRIN lens implant sites in PV-cre mice (N=4), example Ca^2+^ responses of PV BLA interneurons to CS+ and US presentations during fear conditioning and percentage of cells with significantly increased or decreased Ca^2+^ responses during stimulus presentations (n=46). **(G)** Example image of GCaMP6 expression in SOM interneurons in the BLA of SOM-cre mice. Scale bar, 20 μm. **(H)** Left to right, GRIN lens implant sites in SOM-cre mice (N=5), example Ca^2+^ responses to CS+ and US presentations during fear conditioning and percentage of SOM interneurons with significantly increased or decreased Ca^2+^ responses during stimulus presentations (n=152). **(I)** Fraction of US responsive VIP, PV and SOM BLA interneurons averaged across mice (Here and following: VIP, N=7 mice; PV, N=4; SOM, N=5; US excited: Kruskal-Wallis test, H=9.759, P<0.01; Dunn’s multiple comparisons test, VIP vs SOM, P<0.05, PV vs SOM, P<0.05; US inhibited: Kruskal-Wallis test, H=8.639, P<0.01; Dunn’s multiple comparisons test, VIP vs SOM, P<0.05). **(J)** Fraction of CS+ responsive VIP, PV and SOM BLA interneurons averaged across mice (CS+ inhibited: Kruskal-Wallis test, H=7.815, P<0.05; Dunn’s multiple comparisons test, VIP vs SOM, P<0.05). **(K)** Fraction of CS responsive VIP, PV and SOM BLA interneurons averaged across mice. Box-whisker plots show median values and 25^th^/75^th^ percentiles with 10^th^ to 90^th^ percentile whiskers, dots additionally indicate the mean. Bar graphs are mean and s.e.m. * P<0.05. Details of statistical analysis are listed in Table S3.

**Figure S2.**
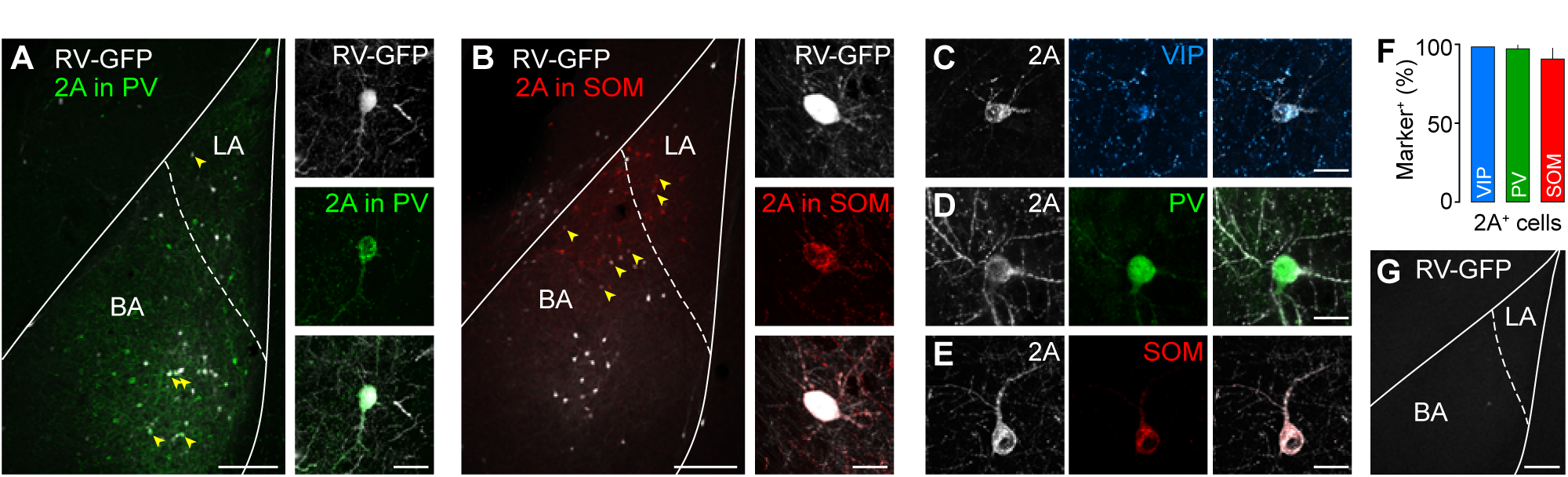
Monosynaptic rabies tracing from VIP, PV and SOM interneurons in the BLA. **(A)** Representative example image of 2A peptide and rabies-GFP (RV-GFP) expression in PV interneurons in the BLA of PV-cre mice. Yellow arrowheads point to identified starter cells expressing both TVA950-2A-CVS11G construct and RV-GFP. LA, lateral amygdala; BA, basal amygdala. Scale bar, 200 μm. High magnification image depicts an example starter cell. Scale bar, 20 μm. **(B)** Co-expression of 2A peptide and RV-GFP in SOM interneurons in the BLA of SOM-cre mice. Yellow arrowheads point to identified starter cells expressing both TVA950-2A-CVS11G and RV-GFP. Scale bar, 200 μm and 20 μm (high magnification). **(C-F)** Specificity of TVA950-2A-CVS11G expression. Example images show co-expression of TVA950-2A-CVS11G and (C) VIP, (D) PV and (E) SOM in VIP-cre, PV-cre and SOM-cre mice, respectively. Scale bars, 20 μm. **(F)** Quantification of co-localization of 2A peptide with interneuron markers detected by immunohistochemistry (VIP, N=1 mouse; PV, N=2; SOM, N=2). **(G)** Representative image illustrating absence of RV-GFP expression in the BLA without preceding TVA950-2A-CVS11G injection. Scale bar, 200 μm. Data is presented as mean and s.e.m.

**Figure S3.**
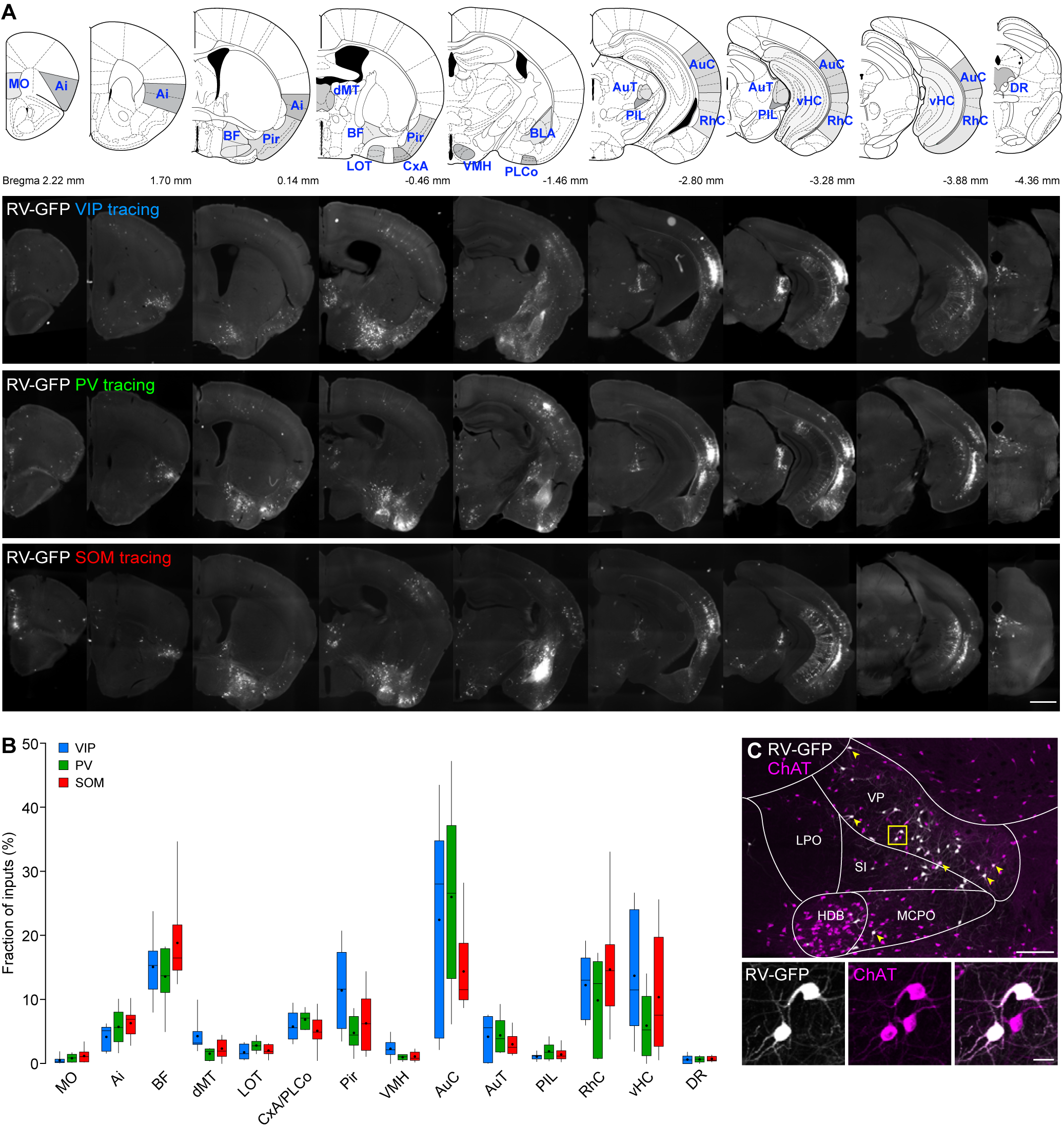
Monosynaptic inputs to VIP, PV and SOM interneurons in the BLA. Serial reconstruction of representative example mouse brains depicting monosynaptic inputs to VIP (top), PV (middle) and SOM (bottom) BLA interneurons. Corresponding injection sites are shown in Figure 2 (VIP, mouse #8 in Figure 2 heatmap, 49% LA starter cells) and Figure S2 (PV, mouse #6, 58% LA; SOM, mouse #8, 38% LA). Top row displays matching mouse brain atlas planes. MO, medial orbital cortex; Ai, agranular insular cortex; BF, basal forebrain; Pir, piriform cortex; dMT, dorsal midline thalamic nuclei; LOT, nucleus of the lateral olfactory tract; CxA, cortex-amygdala transition zone; VMH, ventromedial hypothalamus; PLCo, posterolateral cortical amygdaloid nucleus; BLA, basolateral amygdala; AuT, auditory thalamus; PIL, posterior intralaminar thalamus; AuC, auditory cortices; RhC, rhinal cortices; vHC, ventral hippocampus; DR, dorsal raphe nucleus. Scale bar, 1 mm.Fraction of inputs over total input numbers for each identified brain area projecting to VIP, PV and SOM BLA interneurons (VIP, N=8 mice; PV, N=6; SOM, N=9). **(C)** A subset of basal forebrain presynaptic inputs to VIP BLA interneurons expresses choline acetyltransferase (ChAT; 19.7±4.6%, N=3). LPO, lateral preoptic area; V P, ventral pallidum; SI, substantia innominata, basal part; HDB, nucleus of the horizontal limb of the diagonal band; MCPO, magnocellular preoptic nucleus. Scale bars, 200 μm and 20 μm (high magnification). Data is presented as median values and 25^th^/75^th^ percentiles with 10^th^ to 90^th^ percentile whiskers, dots additionally indicate the mean. Details of rabies tracing analysis are specified in Table S1.

**Figure S4.**
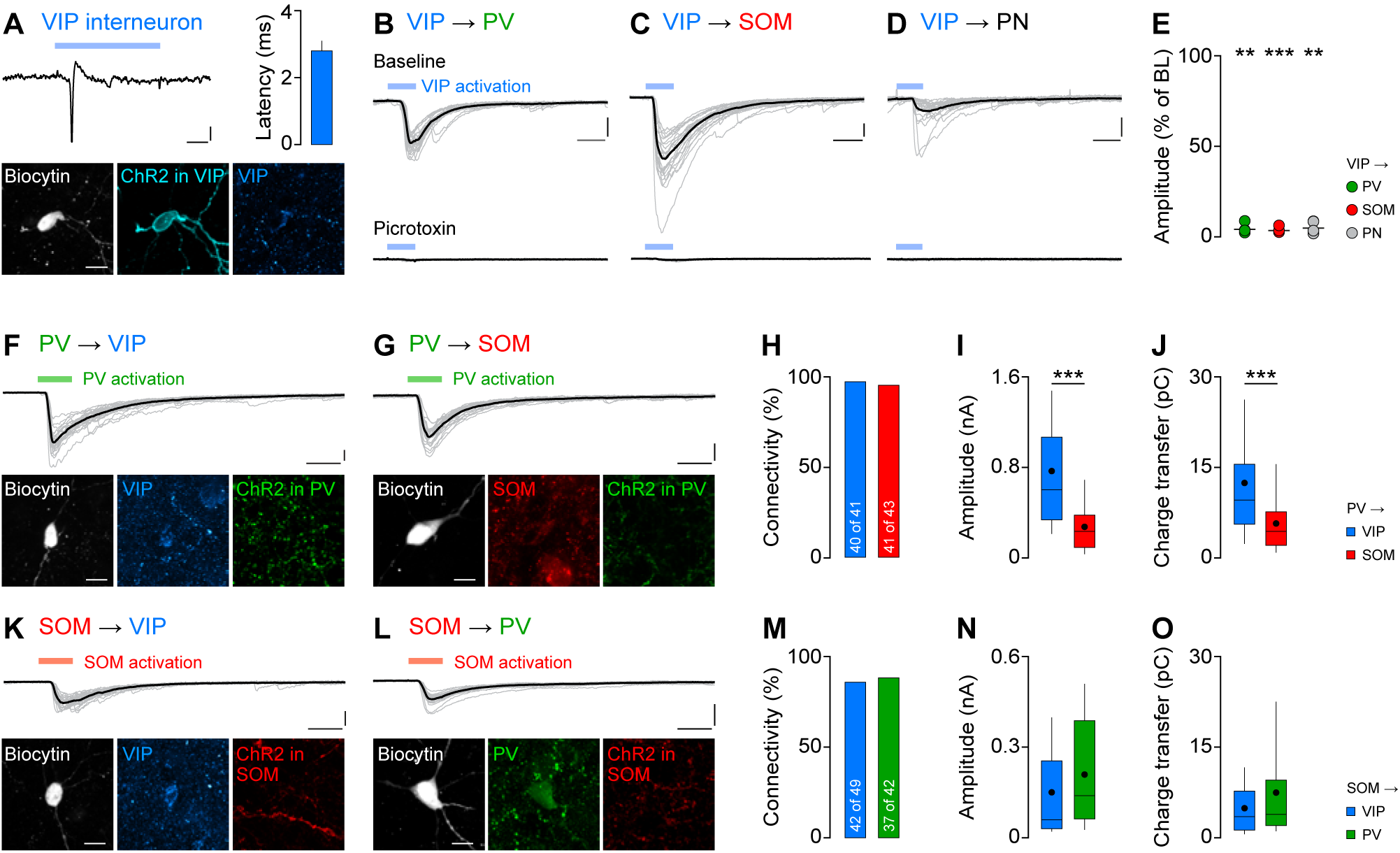
Interconnectivity of BLA interneuron subtypes. **(A)** ChR2-EYFP in VIP BLA interneurons. Top left, example recording of action potential generation by ChR2 activation with blue light in cell-attached mode. Scale bars, 20 pA, 2 ms. Top right, average latency to light-evoked action potentials by ChR2 activation (n=7 cells). Bottom, confocal image of the example cell expressing ChR2 filled with biocytin during whole-cell recordings to confirm VIP expression. Scale bar, 20 μm. **(B-D)** Example traces of IPSCs evoked by VIP BLA network photostimulation before and after application of the GABA_A_-receptor antagonist picrotoxin in (B) PV interneurons, (C) SOM interneurons and (D) PNs of the BLA. Scale bars, 100 pA, 10 ms. **(E)** The amplitude of light-evoked IPSCs is significantly reduced by picrotoxin in PV (ratio paired t-test, P<0.01, n=4) and SOM (ratio paired t-test, P<0.001, n=4) interneurons as well as PNs (ratio paired t-test, P<0.01, n=4). **(F)** Top, example recording from a VIP BLA interneuron receiving short-latency inhibitory inputs upon PV BLA interneuron network activation with ChR2 (green bar). Scale bars, 100 pA, 10 ms. Bottom, corresponding confocal image confirming VIP expression in the biocytin-filled cell. Scale bar, 20 μm. **(G)** Top, example traces of IPSCs in a SOM BLA interneuron upon brief PV BLA interneuron network activation with ChR2 (green bar). Scale bars, 100 pA, 10 ms. Bottom, corresponding confocal image confirming SOM expression. Scale bar, 20 μm. **(H)** High connectivity from PV BLA interneurons to VIP and SOM interneurons (VIP, 97.6%, 40 of 41 cells from N=3 mice; SOM, 95.3%, 41 of 43 cells from N=3 mice). **(I)** IPSC amplitudes are higher in VIP BLA interneurons compared to SOM interneurons (Mann-Whitney U test, P<0.0001; VIP, n=40; SOM, n=41). **(J)** Charge transfer in VIP BLA interneurons is larger compared to SOM interneurons (Mann-Whitney U test, P<0.001; VIP, n=40; SOM, n=41). **(K)** Top, example traces from a VIP BLA interneuron receiving short-latency inhibitory inputs by brief SOM BLA interneuron network activation with ChR2 (red bar). Scale bars, 100 pA, 10 ms. Bottom, corresponding confocal image confirming VIP expression in the recorded cell. Scale bar, 20 μm. **(L)** Top, recording of IPSCs in a PV BLA interneuron upon SOM BLA interneuron network activation with ChR2 (red bar). Scale bars, 100 pA, 10 ms. Bottom, corresponding confocal image confirming PV expression. Scale bar, 20 μm. **(M)** High connectivity from SOM BLA interneurons to VIP and SOM interneurons (VIP, 85.7%, 42 of 49 cells from N=4 mice; PV, 88.1%, 37 of 42 cells from N=3 mice). **(N-O)** Neither IPSC (N) amplitude nor (O) charge transfer upon SOM BLA network photostimulation are different between VIP and PV interneurons. Individual traces from one cell are gray, corresponding average IPSC is shown in black in panels (B-D), (F-G), (K-L). Bar graph in panel (A) represents mean and s.e.m. Dots in panel (E) represent individual data points, horizontal lines additionally indicate the mean. Box-whisker plots show median values and 25^th^/75^th^ percentiles with 10^th^ to 90^th^ percentile whiskers, dots additionally indicate the mean. ** P<0.01, *** P<0.001. Further details of slice electrophysiology analysis are summarized in Table S2. All details of statistical analysis are listed in Table S3.

**Figure S5.**
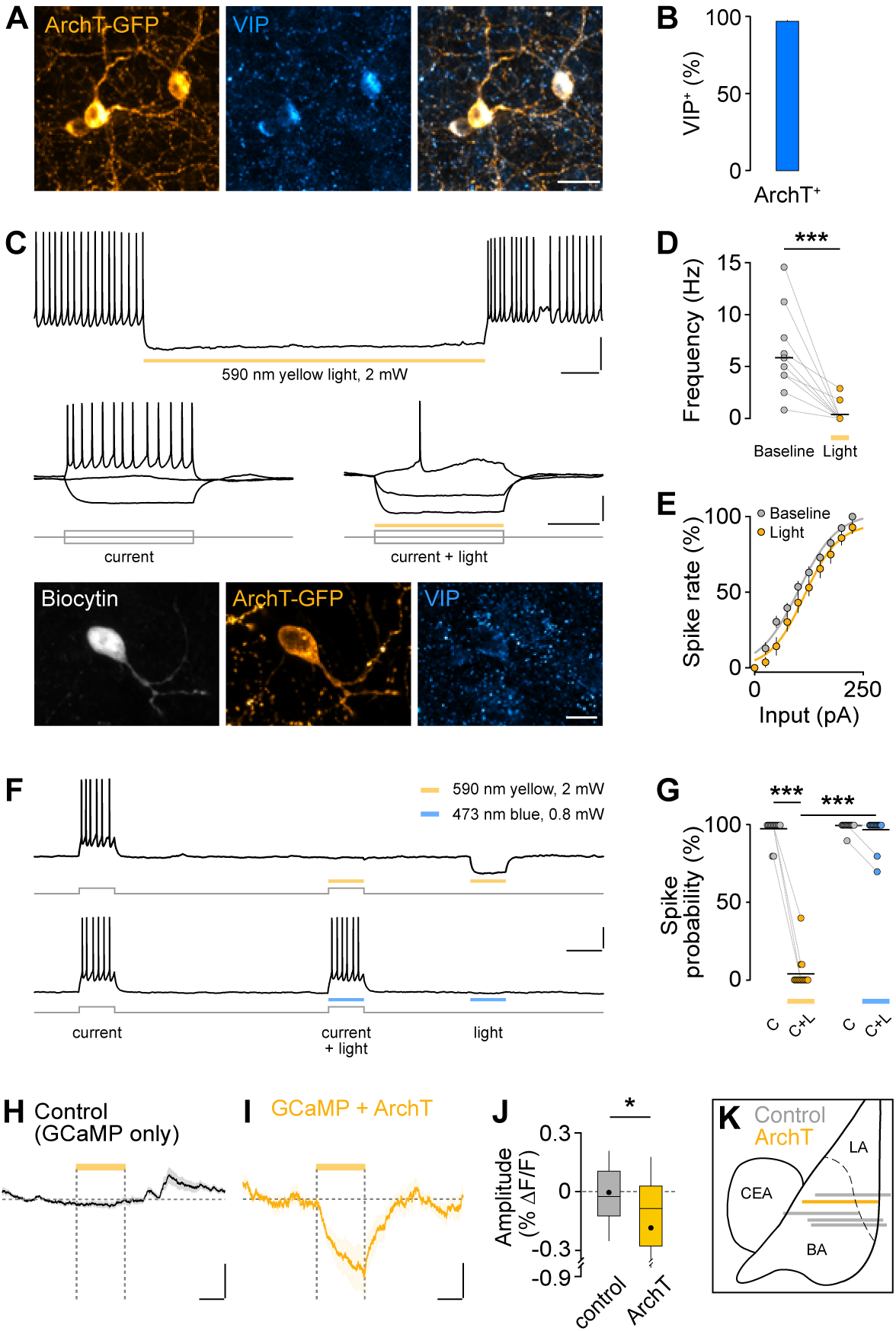
ArchT expression in VIP BLA interneurons. **(A)** Representative example image of ArchT-GFP expression in VIP interneurons in the BLA of VIP-cre mice. Scale bar, 20 μm. **(B)** Quantification of co-localization of ArchT-GFP expression with VIP detected by immunohistochemistry (N=3 mice). **(C)** Representative patch-clamp recording of an ArchT-GFP expressing VIP BLA interneuron. Top, suppression of spontaneous action potential generation by 4.5 s yellow light. Scale bars, 20 mV, 500 ms. Middle, ArchT activation with yellow light diminishes action potentials evoked by depolarizing current steps (−50 pA, 0 pA, and +50 pA current injections while holding the cell at −60 mV). Scale bars, 20 mV, 200 ms. Bottom, confocal image of the same ArchT-GFP+ cell filled with biocytin during whole-cell recordings to confirm VIP expression. Scale bar, 10 μm. **(D)** Spontaneous action potentials are reliably inhibited by application of yellow light (Wilcoxon matched-pairs signed rank test, P<0.001, n=11 cells). **(E)** ArchT activation decreases excitability of VIP BLA interneurons. Spike rate was normalized to the maximum frequency in baseline condition for each cell. Sigmoidal curve fitting reveals a significant shift to the right of input-output curves with ArchT activation (15.7 pA shift; I_half_ baseline 99.9 pA, I_half_ light 115.4 pA; paired t-test, P<0.05; n=12 cells) and decreased gain (I_half_ slope baseline 43.5%/pA, light 36.6%/pA; paired t-test, P<0.05; n=12 cells) without affecting maximum output. **(F)** Representative example traces from a VIP BLA interneuron expressing ArchT-tdTomato, demonstrating reliable spike suppression with yellow, but not blue light. Further, only yellow but not blue light activates ArchT at a holding potential of −60 mV, leading to membrane potential hyperpolarization. Scale bars, 20 mV, 500 ms. **(G)** Yellow light significantly decreases spike probability in VIP BLA interneurons, while blue light has no effect (Friedman test, FM=40.71, P<0.001; Dunn’s multiple comparisons test, BL yellow vs yellow light, P<0.001, yellow light vs blue light, P<0.001; n=15 cells). Note that blue light used for nVoke imaging experiments was further of shorter wavelength and lower intensity (448 nm, 0.4-0.7 mW) compared to slice electrophysiology to exclude unwanted cross-excitation of ArchT. **(H)** Yellow light (yellow line, 590 nm, 12 mW, 20 s) does not affect Ca^2+^ fluorescence in VIP BLA interneurons expressing GCaMP6 (n=95 cells from N=3 mice, trace represents mean and s.e.m.). Scale bars, 0.05% ΔF/F, 10 s. **(I)** Yellow light induces a decrease in Ca^2+^ fluorescence in VIP BLA expressing GCaMP6 and ArchT-tdTomato (n=32 from N=1 mouse). Scale bars, 0.05% ΔF/F, 10 s. **(J)** Average amplitude during yellow light application (20 s) is significantly different between GCaMP6 only controls and VIP interneurons expressing GCaMP6 with ArchT (Mann-Whitney U test, P<0.05; control, n=95; ArchT, n=32). **(K)** Schematic illustrating reconstructed implant sites of GRIN lenses within the BLA for VIP nVoke experiments shown in Figure 5 matched to a mouse brain atlas (gray lines, GCaMP6 in VIP, N=3 mice; yellow line, GCaMP6 and ArchT in VIP, N=1). LA, lateral amygdala; BA, basal amygdala; CEA, central amygdala. Connected dots in panel d and g represent individual paired data points, horizontal lines additionally indicate the mean. Box-whisker plot shows median values and 25^th^/75^th^ percentiles with 10^th^ to 90^th^ percentile whiskers, dots additionally indicate the mean. All other data is presented as mean and s.e.m. * P<0.05, *** P<0.001. All details of statistical analysis are listed in Table S3.

**Figure S6.**
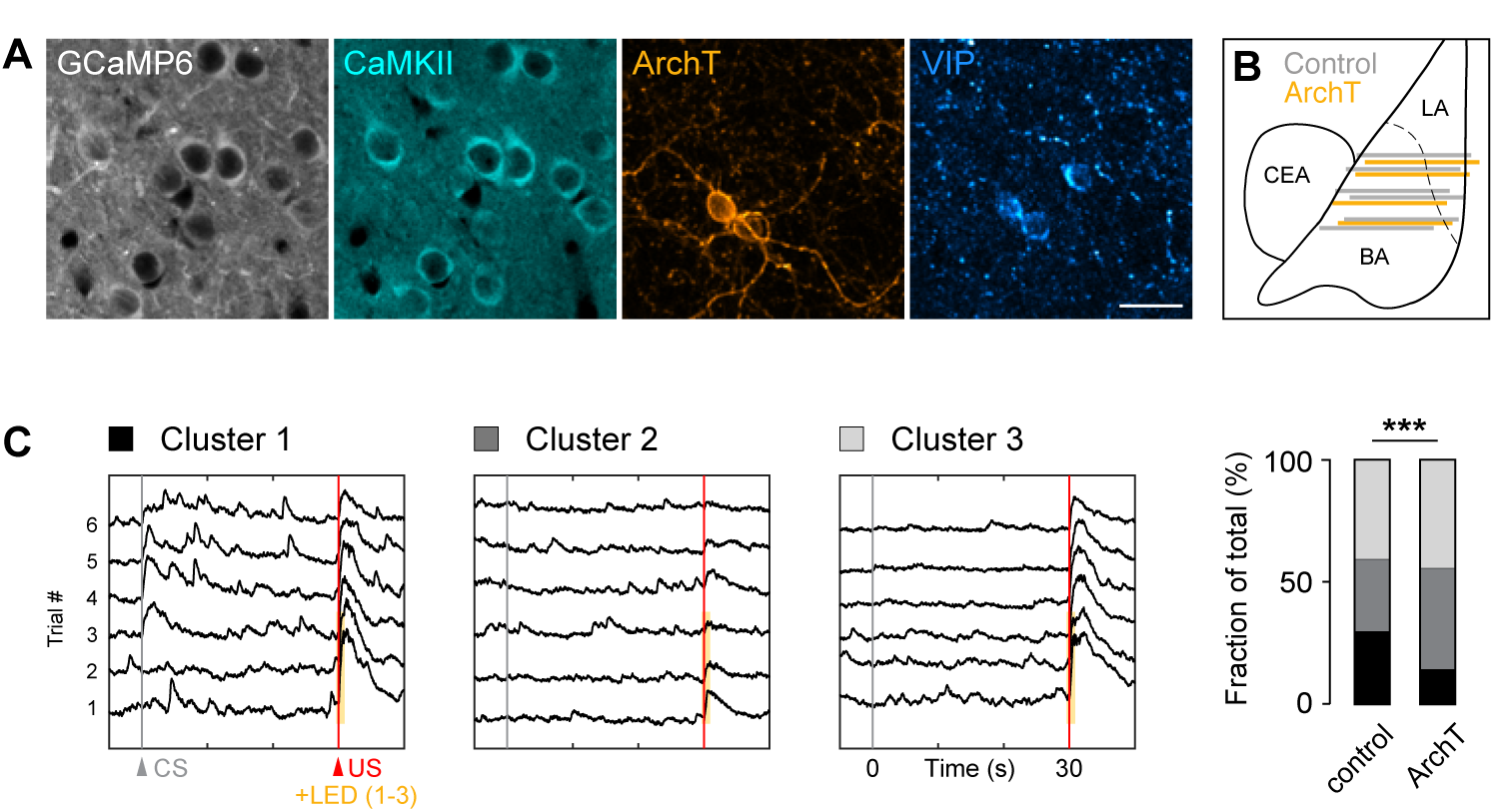
Combined deep brain calcium imaging and optogenetic manipulation. **(A)** Representative example image showing concomitant expression of CaMKII-GCaMP6 and cre-dependent ArchT-tdTomato in the BLA of VIP-cre mice. Immunohistochemical counterstaining against CaMKII and VIP confirms specificity of viral constructs. Scale bar, 20 μm. **(B)** Implant sites of GRIN lenses within the BLA for CaMKII nVoke experiments shown in figure 5 (gray lines, CaMKII-GCaMP6 with tdTomato in VIP, N=6; yellow lines, CaMKII-GCaMP6 with ArchT in VIP, N=4). LA, lateral amygdala; BA, basal amygdala; CEA, central amygdala. **(C)** Average CS and US responses for all pairings for cells clustered based on their CS activity pattern during the last three trials illustrating CS responsive PNs (CS-up pattern, Cluster 1, n=132 cells) or CS non-responsive PNs (Cluster 2: active during both baseline and CS, n=184; Cluster 3: showing no activity during baseline and CS, n=229) from both control and VIP-ArchT mice. Inhibition of VIP interneurons during the US with ArchT significantly reduces changes CS activity patterns in BLA PNs (Pearson’s X^2^ test, P<0.0001; control, n=349; ArchT, n=196). *** P<0.001.

**Figure S7.**
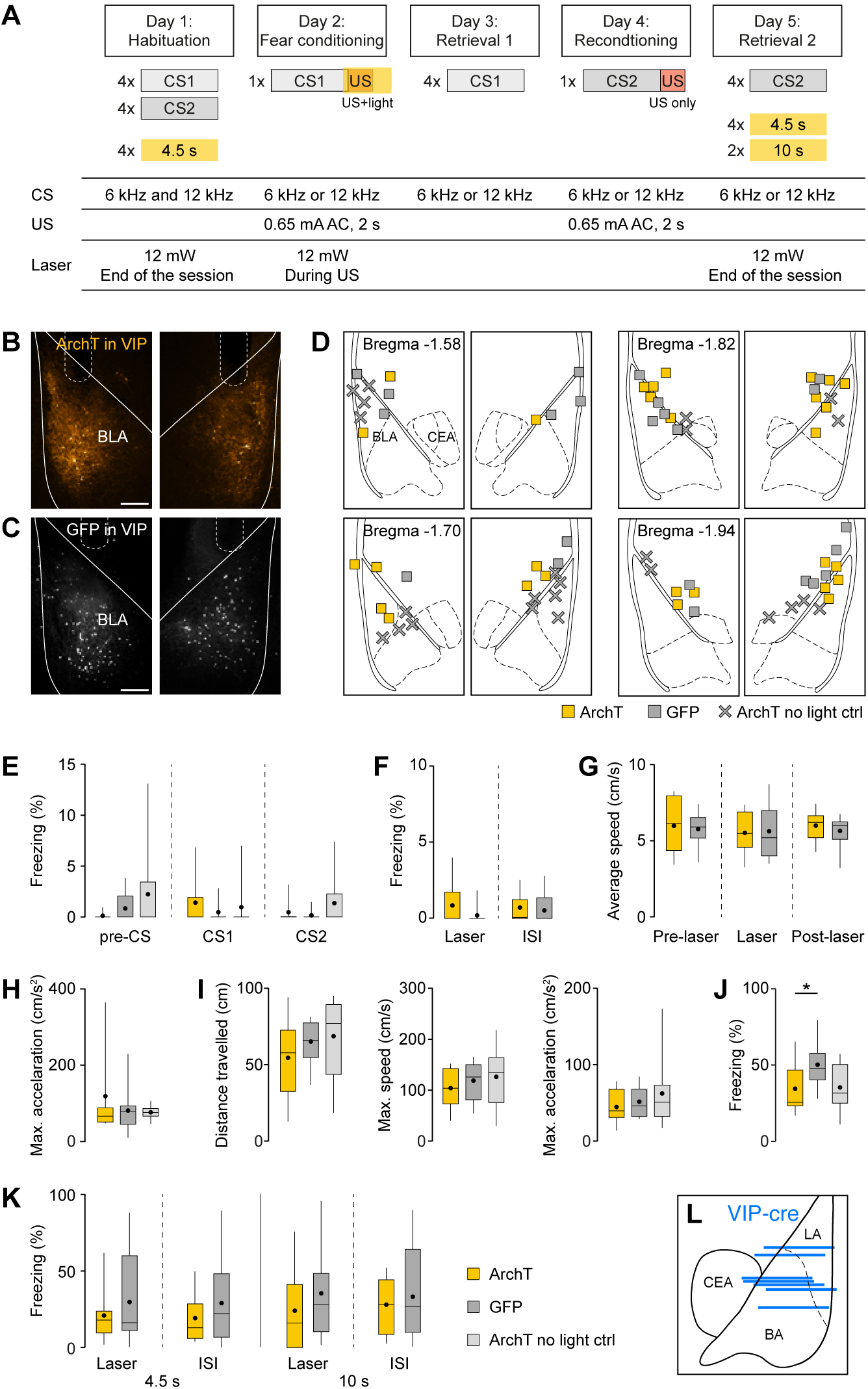
Optogenetic manipulation of VIP BLA interneurons during auditory fear conditioning. **(A)** Schematic illustrating the entire 5-day behavioral paradigm used for optogenetic loss-of-function experiments, including details about CS, US and yellow light pattern applied. **(B-C)** Representative example images of bilateral expression of (B) ArchT-GFP and (C) GFP in VIP interneurons in the BLA of VIP-cre mice with corresponding optical fiber placement (dashed lines). Scale bar, 200 μm. **(D)** Position of optical fiber tips in all mice included in optogenetic experiments matched to a mouse brain atlas. Here and following: ArchT, N=14 mice; GFP, N=11; ArchT no light ctrl, N=12. **(E)** CS presentations on habituation day do not induce freezing in naïve mice. **(F)** Optogenetic inhibition of VIP BLA interneurons has no effect on freezing during or after yellow light stimulus presentation in naïve mice (ISI, inter-stimulus interval). **(G)** Similarly, light stimulation during the habituation session does not affect running speed in either of the light-treated groups. **(H)** Maximum acceleration during the aversive US during fear conditioning. **(I)** Left to right, maximum acceleration, maximum speed and distance travelled during the aversive US during reconditioning. **(J)** Minor differences in post-shock freezing between the ArchT group and GFP controls (Kruskal-Wallis test, H=6.437, P<0.05; Dunn’s multiple comparisons test, ArchT vs GFP, P<0.05). **(K)** Optogenetic inhibition of VIP BLA interneurons for 4.5 s or 10 s at the end of the retrieval 2 session does not affect freezing behavior. **(L)** Implant sites of GRIN lenses (blue lines, N=7) within the BLA of VIP-cre mice for repeated fear conditioning experiments shown in Figure 6. Box-whisker plots show median values and 25^th^/75^th^ percentiles with 10^th^ to 90^th^ percentile whiskers, dots additionally indicate the mean. Bar graphs present mean and s.e.m. * P<0.05. All details of statistical analysis are listed in Table S3.

**Table S1.** Overview of monosynaptic rabies tracing results. Starter cell number and percentage of LA starter cells are presented as mean with s.e.m. and were not significantly different between the three groups (one-way ANOVA). Presynaptic cell number, fraction of inputs and convergence index are shown as median values with 25^th^/75^th^ percentiles. For abbreviations, see Figure S3 or methods section.

|  | VIP BLA tracing | PV BLA tracing | SOM BLA tracing |
| --- | --- | --- | --- |
| **Sample size** | N=8 mice | N=6 | N=9 |
| **Starter cell #** | 121±26 | 57±9 | 72±17 |
| **LA starter (%)** | 53.7±11.9 | 57.6±15.5 | 58.2±8.8 |
| **Presynaptic cell #** |  |  |  |
| MO | 14 (10/34) | 38 (15/83) | 35 (12/145) |
| Ai | 198 (96/250) | 244 (121/446) | 246 (144/508) |
| BF | 598 (453/870) | 579 (450/833) | 780 (644/1186) |
| dMT | 161 (116/265) | 59 (29/100) | 189 (44/222) |
| LOT | 84 (29/127) | 116 (103/136) | 95 (74/116) |
| CxA/PLCo | 273 (237/384) | 238 (199/434) | 185 (141/532) |
| Pir | 520 (218/702) | 177 (121/393) | 237 (114/448) |
| VMH | 111 (61/150) | 34 (33/82) | 67 (24/85) |
| AuC | 1180 (257/1512) | 1312 (973/1638) | 576 (303/832) |
| AuT | 248 (16/301) | 154 (116/263) | 81 (69/186) |
| PIL | 46 (35/67) | 87 (33/129) | 53 (36/85) |
| RhC | 473 (356/769) | 453 (123/897) | 691 (355/1009) |
| vHC | 701 (352/1022) | 153 (82/641) | 432 (95/855) |
| DR | 21 (0/57) | 28 (15/54) | 23 (14/59) |
| **Fraction of inputs (%)** |  |  |  |
| MO | 0.34 (0.27/0.57) | 0.93 (0.42/1.28) | 1.13 (0.34/1.57) |
| Ai | 5.15 (1.97/5.70) | 5.65 (4.14/7.25) | 6.92 (5.07/7.06) |
| BF | 15.34 (11.83/17.31) | 13.74 (13.22/16.91) | 16.48 (15.63/18.17) |
| dMT | 3.23 (3.02/4.98) | 1.92 (0.83/2.25) | 1.99 (1.61/3.46) |
| LOT | 1.57 (0.86/2.56) | 2.79 (2.40/3.09) | 1.91 (1.58/2.87) |
| CxA/PLCo | 5.61 (4.14/6.70) | 7.13 (5.77/7.55) | 4.95 (3.97/6.26) |
| Pir | 11.65 (6.76/14.41) | 4.48 (3.61/6.55) | 6.23 (2.30/8.34) |
| VMH | 2.17 (1.56/3.08) | 1.08 (0.78/1.37) | 0.84 (0.74/1.71) |
| AuC | 28.06 (5.26/33.77) | 26.60 (16.60/33.81) | 11.56 (10.16/18.14) |
| AuT | 5.61 (0.33/6.97) | 3.92 (2.09/5.71) | 2.59 (1.81/4.19) |
| PIL | 1.22 (0.81/1.29) | 1.76 (0.95/2.39) | 1.19 (0.84/1.62) |
| RhC | 13.04 (7.30/16.23) | 12.51 (3.17/15.38) | 14.53 (9.27/17.27) |
| vHC | 11.53 (7.71/20.52) | 5.27 (1.87/9.02) | 7.57 (2.73/19.42) |
| DR | 0.60 (0.00/1.13) | 0.70 (0.39/1.01) | 0.55 (0.46/1.17) |
| **Convergence index** |  |  |  |
| MO | 0.17 (0.08/0.36) | 0.90 (0.35/0.99) | 0.50 (0.31/1.17) |
| Ai | 2.98 (0.79/3.27) | 4.47 (3.21/6.62) | 4.80 (3.23/5.93) |
| BF | 8.44 (3.58/9.64) | 12.82 (11.18/13.52) | 13.47 (9.88/16.52) |
| dMT | 2.03 (1.38/2.67) | 1.50 (0.59/1.85) | 1.47 (1.31/2.00) |
| LOT | 0.64 (0.43/1.27) | 2.42 (1.83/2.66) | 1.33 (0.97/2.56) |
| CxA/PLCo | 2.67 (2.04/3.84) | 5.80 (4.33/7.17) | 3.67 (2.52/5.93) |
| Pir | 5.04 (2.07/8.93) | 3.57 (2.47/6.73) | 3.73 (2.53/7.85) |
| VMH | 0.89 (0.70/1.53) | 0.69 (0.69/1.22) | 0.80 (0.49/1.07) |
| AuC | 8.26 (3.36/14.33) | 19.33 (17.78/29.33) | 7.77 (7.27/11.96) |
| AuT | 1.73 (0.22/2.77) | 2.82 (1.86/5.19) | 1.90 (1.18/2.88) |
| PIL | 0.53 (0.33/0.61) | 1.45 (0.69/2.17) | 0.80 (0.53/1.62) |
| RhC | 5.78 (3.08/9.66) | 11.09 (2.79/14.13) | 9.10 (6.50/9.38) |
| vHC | 5.89 (2.97/9.74) | 4.25 (1.63/7.61) | 7.13 (2.44/10.06) |
| DR | 0.31 (0.00/0.62) | 0.72 (0.31/1.00) | 0.47 (0.36/0.77) |

**Table S2.** Overview of connectivity parameters between BLA neuronal subtypes. Connectivity was compared using a Pearson’s X^2^ test with Fisher’s exact post-hoc test. All other statistics were analyzed using a Kruskal-Wallis test and post-hoc Dunn’s multiple comparisons, data are median values and 25^th^/75^th^ percentiles.

| Chr2 in VIP BLA | PV (n=41) | SOM (n=34) | PN (n=22) | Statistics | Post-hoc |
| --- | --- | --- | --- | --- | --- |
| Connectivity | 91.7 %, n=55 of 60 from N=7 mice | 100 %, n=46 of 46 from N=5 mice | 48.6 %, n=35 of 72 from N=13 mice | $P<0.0001$ | PV vs SOM ns<br>PV vs PN $P<0.0001$<br>SOM vs PN $P<0.0001$ |
| Amplitude (pA) | 142.2 (51.6/316.7) | 227.1 (113.1/523.6) | 97.1 (26.7/17.6) | $H=9.162$<br>$P<0.05$ | PV vs SOM ns<br>PV vs PN ns<br>SOM vs PN $P<0.001$ |
| Latency to peak (ms) | 6.7 (5.8/8.1) | 7.15 (5.7/8.7) | 6.8 (5.3/8.5) | ns | - |
| Rise time (10-90 %, ms) | 1.7 (1.3/2.3) | 2.0 (1.4/2.7) | 1.7 (1.3/3.0) | ns | - |
| Decay time (100-37 %, ms) | 7.0 (5.8/8.9) | 16.0 (13.5/17.4) | 11.1 (8.1/14.0) | $H=47.78$<br>$P<0.0001$ | PV vs SOM $P<0.0001$<br>PV vs PN $P<0.01$<br>SOM vs PN $P<0.01$ |
| Charge transfer (pC) | 1.6 (0.7/3.8) | 4.6 (2.3/11.8) | 1.4 (0.7/3.0) | $H=16.85$<br>$P<0.001$ | PV vs SOM $P<0.01$<br>PV vs PN ns<br>SOM vs PN $P<0.01$ |
| Capacitance (pF) | 35.7 (30.6/45.5) | 53.7 (42.3/65.0) | 126.3 (102.6/139.0) | $H=62.32$<br>$P<0.0001$ | PV vs SOM $P<0.01$<br>PV vs PN $P<0.001$<br>SOM vs PN $P<0.001$ |
| Conductance (pA/mV) | 3.0 (1.2/8.0) (n=22) | 7.1 (3.7/13.7) (n=26) | 2.3 (1.2/4.1) (n=11) | $H=7.797$<br>$P<0.05$ | PV vs SOM ns<br>PV vs PN ns<br>SOM vs PN $P<0.05$ |

| Chr2 in PV BLA | VIP (n=40) | SOM (n=41) | PN (n=34) | Statistics | Post-hoc |
| --- | --- | --- | --- | --- | --- |
| Connectivity | 97.6 %, n=40 of 41 from N=3 mice | 95.3 %, n=41 of 43 from N=3 mice | 97.1 %, n=34 of 35 from N=6 mice | ns | - |
| Amplitude (pA) | 598.4 (331.7/1066.0) | 229.0 (87.0/372.6) | 1429.0 (816.1/2876.0) | $H=50.73$<br>$P<0.0001$ | VIP vs SOM $P<0.0001$<br>VIP vs PN $P<0.05$<br>SOM vs PN $P<0.0001$ |
| Latency to peak (ms) | 5.9 (4.9/6.7) | 7.6 (6.1/9.1) | 6.4 (5.5/8.1) | $H=21.32$<br>$P<0.0001$ | VIP vs SOM $P<0.0001$<br>VIP vs PN $P<0.05$<br>SOM vs PN ns |
| Rise time (10-90 %, ms) | 1.5 (1.1/1.9) | 2.7 (1.8/3.5) | 1.7 81.4/3.0) | $H=23.8$<br>$P<0.0001$ | VIP vs SOM $P<0.0001$<br>VIP vs PN ns<br>SOM vs PN $P<0.05$ |
| Decay time (100-37 %, ms) | 9.4 (8.5/11.2) | 13.6 (11.0/16.1) | 15.2 (13.0/19.9) | $H=45.22$<br>$P<0.0001$ | VIP vs SOM $P<0.0001$<br>VIP vs PN $P<0.0001$<br>SOM vs PN ns |
| Charge transfer (pC) | 9.5 (5.6/15.5) | 4.4 (2.0/7.6) | 33.4 (12.5/74.7) | $H=48.06$<br>$P<0.0001$ | VIP vs SOM $P<0.01$<br>VIP vs PN $P<0.001$<br>SOM vs PN $P<0.0001$ |
| Capacitance (pF) | 26.3 (22.0/30.2) | 45.6 (37.9/54.0) | 106.5 (90.9/117.8) | $H=92.96$<br>$P<0.0001$ | VIP vs SOM $P<0.0001$<br>VIP vs PN $P<0.0001$<br>SOM vs PN $P<0.0001$ |
| Conductance (pA/mV) | 7.7 (5.3/14.0) (n=36) | 3.3 (1.9/5.5) (n=33) | 27.0 (12.4/52.9) (n=33) | $H=43.3$<br>$P<0.0001$ | VIP vs SOM $P<0.01$<br>VIP vs PN $P<0.01$<br>SOM vs PN $P<0.0001$ |

**Table S2. Overview of connectivity parameters between BLA neuronal subtypes.**
| Chr2 in SOM BLA | VIP (n=42) | PV (n=37) | PN (n=33) | Statistics | Post-hoc |
| --- | --- | --- | --- | --- | --- |
| Connectivity | 85.7 %, n=42 of 49 from N=4 mice | 88.1 %, n=37 of 42 from N=3 mice | 100 %, n=33 of 33 from N=7 mice | ns | - |
| Amplitude (pA) | 145.1 (67.6/394.2) | 64.8 (34.8/260.3) | 820.8 (253.8/1116.0) | $H=38.29$<br>$P<0.0001$ | VIP vs PV ns<br>VIP vs PN $P<0.0001$<br>PV vs PN $P<0.0001$ |
| Latency to peak (ms) | 8.2 (7.4/9.9) | 8.4 (7.1/9.7) | 10.1 (7.2/12.0) | ns | - |
| Rise time (10-90 %, ms) | 2.3 (1.6/3.7) | 2.9 (2.1/4.2) | 2.8 (2.0/3.6) | ns | - |
| Decay time (100-37 %, ms) | 12.4 (9.5/15.0) | 26.0 (14.2/39.3) | 22.9 (18.5/26.9) | $H=33.75$<br>$P<0.0001$ | VIP vs PV $P<0.0001$<br>VIP vs PN $P<0.0001$<br>PV vs PN ns |
| Charge transfer (pC) | 3.5 (1.3/7.7) | 3.9 (2.0/9.5) | 31.5 (7.4/43.4) | $H=33.58$<br>$P<0.0001$ | VIP vs PV ns<br>VIP vs PN $P<0.0001$<br>PV vs PN $P<0.0001$ |
| Capacitance (pF) | 24.7 (20.5/29.4) | 35.4 (30.0/43.2) | 105.2 (81.5/120.5) | $H=81.56$<br>$P<0.0001$ | VIP vs PV $P<0.001$<br>VIP vs PN $P<0.0001$<br>PV vs PN $P<0.0001$ |
| Conductance (pA/mV) | 2.5 (0.9/7.0) (n=36) | 1.9 (1.0/4.2) (n=27) | 16.5 (5.6/30.2) (n=33) | $H=37.45$<br>$P<0.0001$ | VIP vs PV ns<br>VIP vs PN $P<0.0001$<br>PV vs PN $P<0.0001$ |

**Table S3.**
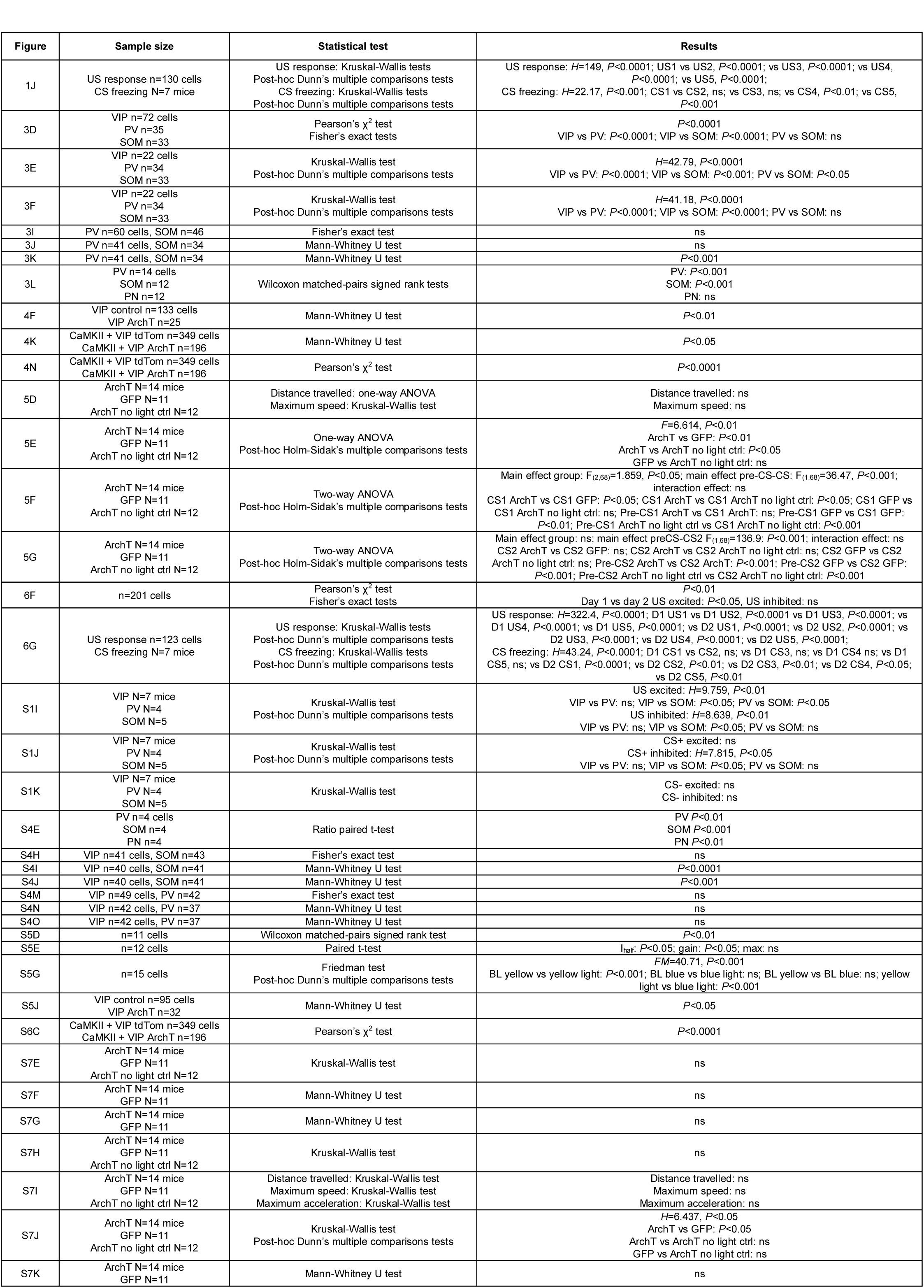
Summary of all statistical analyses for data presented in main and supplementary figures.

